# Human trunk embryoids with patterned anterior-posterior and dorsal-ventral body axes: utility for understanding human development and disease

**DOI:** 10.64898/2025.12.20.695666

**Authors:** Tianming Wu, Hao Yu, Brian S.H. Wong, Kexin Teng, Weiman Xiang, Ling Xu, Jianan Zhang, Angel Y.F. Kam, Ethel S.K. Ng, Joaquim Vong, Jiannan Zhang, Bo Gao, Stephen K.W. Tsui, Stephen Dalton

**Affiliations:** School of Biomedical Sciences, The Chinese University of Hong Kong, Shatin, Hong Kong SAR, China; School of Biomedical Sciences, The University of Hong Kong, Hong Kong SAR, China; Key Laboratory of Bio-resources and Eco-environment of Ministry of Education, College of Life Sciences, Sichuan University, Chengdu, China

**Keywords:** pluripotent stem cells, embryoid, bipotent NMPs, notochord, A-P and D-V axes, human trunk development

## Abstract

Human embryoid models enable mechanistic studies of development and disease. We generated trunk embryoids from human pluripotent stem cells that recapitulate posterior trunk formation at Carnegie stage (CS) 8-10, with patterned anterior-posterior (A-P) and dorsal-ventral (D-V) axes. These self-organizing structures comprise a ventral notochord, dorsal neural tube, floor plate and bilateral somites. Genetic and chemical perturbations of SHH signaling confirmed the notochord’s central role in D-V patterning. Moreover, VANGL1/2 loss-of-function mutations recapitulated mouse phenotypes, including axial truncation and somite segmentation failure. This model enables detailed study of key developmental events that underlie posterior trunk formation and provides a promising platform for human disease modeling.

## Introduction

Understanding the molecular and cellular aspects of peri- and post-implantation human embryogenesis are fundamental for a better understanding of congenital disease, the application of human pluripotent stem cell (hPSC) towards regenerative medicine and potentially, drug validation. Developing hPSC-derived embryoid models offers opportunities to address these issues.^1–3^ Although success has been achieved in developing mouse embryoids that exhibit multi-axial and multi-tissue patterning along the anterior-posterior (A-P), dorsal-ventral (D-V) and left-right (L-R) body axes,^4–6^ attempts to generate multi-axial human trunk models suffer from several limitations.

Initial human trunk models using bipotent neuromesoderm progenitors (bi-NMPs) are uni-axial and comprised of somite-only^7–9^ or neural tube-only^10^ structures. Recently developed coupled models^11–13^ are composed of a neural tube and somites but, are often aberrantly structured (e.g. unilateral somites and twisted neural tube) and fail to establish a D-V body axis. Notably, these models lack a notochord and are heavily biased towards dorsal patterning.^12,13^ In contrast, trunk models that form a notochord-like structure are ventrally-biased and lack a neural tube and somites.^14^ These limitations highlight the need for a human model that supports co-development of the notochord alongside bi-NMPs, together with the establishment of the A-P and D-V body axes.

Temporal signaling by WNT, NODAL, BMP, retinoic acid (RA), FGF and SHH specifies tailbud bi-NMPs and notochord progenitor cells along the A-P and D-V axes.^14–18^ A major challenge is to establish a concerted signaling environment that supports these co-developmental events. We iteratively optimized conditions to generate human trunk embryoid models (hTEMs) from bi-NMPs and notochord progenitors. hTEMs self-organize into notochord, dorsal neural tube, floor plate, and bilateral somites. Validation of this was based on morphological, cellular and molecular criteria using spatial and single-cell transcriptomics, live imaging and scanning electron microscopy (SEM). Comparative analyses showed that hTEMs recapitulate posterior trunk development equivalent to that in human embryos at Carnegie stages (CS) 8-10.

The utility of hTEMs for modeling human development was investigated by perturbing notochord identity and notochord-derived SHH activity. By this approach, the notochord was shown to be a major driving force for D-V axis establishment in the human trunk. Moreover, hTEMs faithfully reproduced human neural tube defects (NTDs), as shown by loss-of-function mutations in the planar cell polarity (PCP) genes, *VANGL1* and *VANGL2*. These findings highlight the versatility of trunk embryoids for understanding human development and congenital disease and points towards additional applications for drug discovery and regenerative medicine.

## Results and Discussion

### Dorsally biased neural tube-somite coupled embryoids

We first aimed to generate topographically coupled embryoids with A-P segmented somites flanking an elongating neural tube (hTEM.v1; Figures 1A and 1B). To achieve this, size-controlled 3D human embryonic stem cell (hESC) spheroids received WNT agonist CHIR99021 (CHIR), FGF2, and inhibitors for BMP (LDN-193189 (LDN)) and NODAL (SB-431542 (SB)) pathways. Resultantly, hTEM.v1 embryoids became asymmetric and by day4, underwent mediolateral narrowing and axial elongation (Figures 1B and S1A). Matrigel and retinal (RAL) promoted axial elongation and somite segmentations at days 4-7 (Figures 1C, S1B, and S1C).^8^ 6-8 pairs of bilaterally positioned somites progressively formed along with gradual extension and closure of the neural tube (Figures 1D, S1D, and Video S1). By day 7, hTEM.v1 reached a mean length of 1857 μm (Figure 1C).

**Figure 1.**
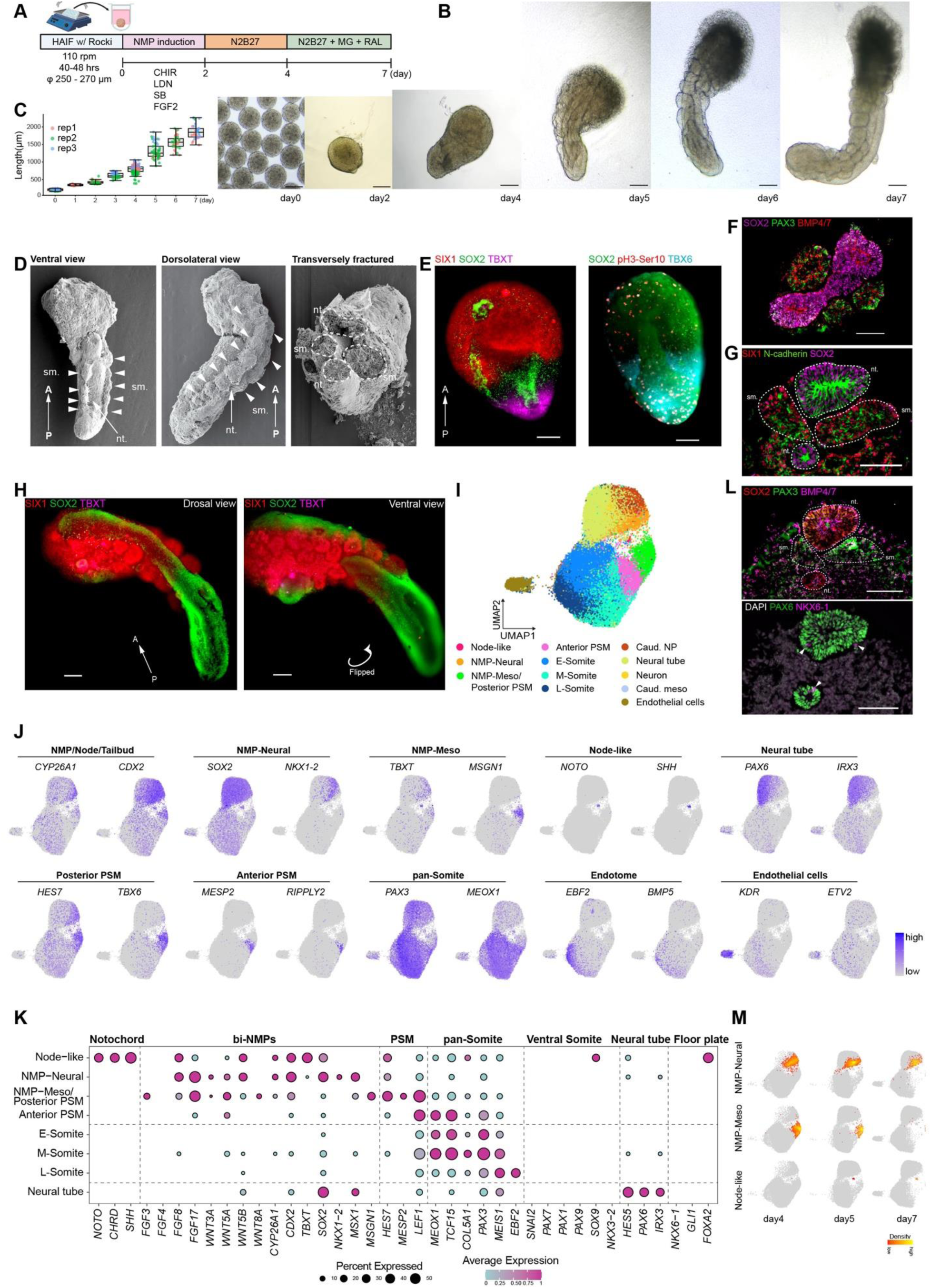
Generation of dorsally biased fates hTEM.v1. (A) Schematic of the hTEM.v1 generation. (B) Representative images of hTEM.v1 from day 0 to day 7. Scale bars, 200 μm. (C) Box plot of embryoid length over time from three independent biological replicates (rep). Each dot represents an individual hTEM.v1 length measurement (n = 9-124 for each time point). (D) SEM snapshots of day-7 hTEM.v1 samples (n = 3) in longitudinal orientation from ventral (left), dorsolateral (middle) and transversely fractured views. nt., neural tube. sm., somite. (E) 3D projections of representative day-4 hTEM.v1 stained for NMPs (TBXT, SOX2), PSM (TBX6), early unsegmented somite (SIX1), and mitotic cells (pH3-Ser10). Scale bars, 100 μm. (F) Immunofluorescence image of longitudinally sectioned day-6 hTEM.v1 showing the neural tube (SOX2) and flanking pairs of somites (PAX3). Scale bar, 100 μm. (G) Immunofluorescence images of a transversely sectioned day-7 hTEM.v1 showing the spatial organization of duplicated, epithelialized (N-cadherin) neural tube structures (SOX2) relative to paired somites (PAX3). (H) 3D projections of day-7 hTEM.v1 from the dorsal (left) and ventral (right) displaying the duplicated neural tubes (SOX2) with flanking somites (SIX1). Caudally positioned NMPs (TBXT) were barely detectable on day 7. Scale bars, 100 μm. (I) UMAP of integrated hTEM.v1 integration from day 4/5/7 (total of 47,908 cells). Cell type annotations are indicated below. (J) UMAP showing the expression of selected cell types from hTEM.v1. (K) Dot plot displaying the expression levels (color) and proportion (dot size) of marker genes for dorsal and ventral fates. Proportions below 5% were omitted. (L) (Top) Both neural tube (SOX2, PAX3/6, BMP4/7) and somite cell (PAX3, BMP4/7) identities were dorsal-biased in day-7 hTEM.v1, (bottom) with few detectable ventral neural tube (NKX6-1, arrowhead) cells. Scale bars, 100 μm. (M) Density plot showing the progress of NMP-neural, NMP-Meso and Node-like cells over time.

Immunostaining of hTEM.v1 (days 2-4) revealed the following features indicative of A-P patterning; (i) polarized expression of TBXT and CDX2 at the posterior end, (ii) mutually exclusive positioning of central neural (SOX2) and mediolateral pre-somitic mesoderm (PSM; TBX6) cells; (iii) A-P symmetry breaking, indicated by anterior somitic (SIX1) cells undergoing an epithelial-to-mesenchymal transition (EMT; N-cadherin) and caudalized tailbud (CDX2) cells (Figures 1E, S1E-S1G). In total, these characteristics are reminiscent of the highly mitotic (pH3-Ser10), A-P patterned human CS8 embryo (Figure 1E).^19,20^ Upon further inspection of hTEM.v1 (days 4-7), SOX2+TBXT-neural tubes were observed on both dorsal and ventral sides along the A-P axis (Figures 1F-1H and S1H). This is coincident with the absence of a SOX2-TBXT+ notochord structure (Figures 1E and S1I). This suggests that although the A-P axis had formed, D-V patterning in hTEM.v1 was not established.

To understand D-V patterning defects in hTEM.v1, single-cell RNA sequencing (scRNA-seq) analysis was performed (days 4/5/7). Clustering and hierarchical transcriptomic profiling identified 12 major cell identities, including two major continuums delineating somitogenesis and neural tube formation (Figures 1I, 1J, S1J, and S1K); (i) the somitic lineage contains NMP-Meso (*TBXT*, *MSGN1*), posterior PSM (*HES7*, *TBX6*), anterior PSM (*MESP2*, *RIPPLY2*) and pan-somite cells (*PAX3*, *SIX1*, comprised of early-, middle- and late-somite (E-, M- and L-somite) subtypes along the time course; (ii) the neural lineage includes NMP-Neural (*SOX2*, *NKX1-2*), caudal neural plate progenitors (Caud. NP; *MSX2*, *SOX2*, *NKX1-2*) and neural tube cells (*PAX6, HES5*).

Overall, due to the absence of ventral somite compartments (*PAX1*, *PAX9*) or ventral neural tube (*NKX6-1*) and floor plate (*FOXA2*), the spatiotemporal expression profile of hTEM.v1 confirmed dorsally-biased cell identities in both the somitic and neural lineages (Figure 1K). Immunostaining of transversely sectioned day-7 hTEM.v1 further confirmed the unrestricted distribution of dorsal BMP4/7 signals and over-expansion of dorsal neural (PAX6) cells, with few to no ventral neural (NKX6-1) cells in the neural tube region (Figure 1L).

Conventionally, bi-NMPs are identified by the co-expression of SOX2 and TBXT,^21–23^ but questions about their molecular identities remain unresolved.^24–26^ Here, we identified distinct NMP-Neural (*SOX2*^high^,*TBXT*^low^) and NMP-Meso (*SOX2*^low^,*TBXT*^high^) subtypes that exhibited contrasting expression patterns (Figures 1J and 1M). Additionally, they expressed differential levels of *FGF3/4/8/17*, *WNT3A/5A/5B/8A*, and *BM2/4/7* (Figure S1L), which are believed to be the driving force for bi-NMP fate bifurcations in human trunk formation.^27^ This suggests hTEM.v1’s potential to resolve the plasticity of bi-NMP bifurcation.

Then, we asked why notochord was absent from hTEM.v1. scRNA-seq combined with *in situ* hybridization chain reaction (HCR) only detected few cells expressing notochord markers *(TBXT, NOTO*, *CHRD*, *SHH),* possibly representing the ventral node (*SHH*) surrounded by PSM (*TBX6*, *HES7*) (Figures 1J, S1M, and S1N).^28^ In human embryos, correct D-V axis specification is initiated by SOX2^high^/TBXT^low^ in the dorsal neural plate and SOX2^low^/TBXT^high^ in the ventral notochordal plate.^20,27^ However, in day-4 hTEM.v1, TBXT protein was restricted to the tailbud and was undetectable by day 7, while SOX2 expression extended to both dorsal and ventral tube structures (Figures 1E, 1G, and S1I). The SOX2 over-expansion and neural tube duplication phenotype in hTEM.v1 coincides with previously reported NTDs caused by mutations/misregulation of Tbxt in mouse notochord progenitors.^29,30^ We, therefore, attributed the neural tube duplication from hTEM.v1 to the misregulation of *TBXT* and consequently, the inability to specify notochord cells.

Overall, bi-NMP-derived hTEM.v1 with spatially coupled neural tube structures and flanking somite segments were generated. Notochord is well-known for its role in D-V patterning of neural tube and somitic cells in the embryo.^31–33^ The absence of a notochord and SHH signaling along the A-P axis in hTEM.v1 explains the heavy dorsalization observed. Moreover, it is noteworthy that we observed a subset of somitic cells resembling the endotome (*EBF2*) (Figure 1J), a poorly understood somite compartment that contributes to the dorsal aorta, endothelium and hematopoietic stem cells.^13,34,35^ The anteriorly positioned EBF2:mScarlet signal coincided with vascular endothelial cells (SOX17, VE-cadherin) at the somite periphery (Figure S1O and Video S1). These findings indicate that hTEM-based embryoids have utility for studying more advanced events related to later stages of gastrulation in humans, such as somitogenesis, neural tube morphogenesis and somite-derived vasculogenesis.^27^

### Exogenous SHH activation induces ventral fates in trunk embryoids

As notochord-derived SHH signaling is key to ventral patterning,^33^ we postulated that SHH activation could establish a D-V axis and rescue the dorsal-biased defects in hTEM.v1. To test this, day-4 hTEM.v1 embryoids were exposed to varied durations and concentrations of Smoothened agonist (SAG) (Figure S2A). SAG-treated hTEM.v1 exhibited elevated transcripts (e.g., ventral neural tube; *NKX6-1*. Sclerotome; *PAX1*/*9*) of ventral identities by day 7, in a dose- and time-dependent manner (Figure S2B). A pulse of 100 nM SAG was chosen to generate hTEM.v2, because this condition established D-V patterning in neural and somitic cells without affecting morphology or axial elongation (Figures 2A-2D and S2B). Immunostaining of day-7 hTEM.v2 confirmed the emergence of ventral (NKX6-1) neural cells in both neural tubes along the A-P axis (Figure 2E). Notably, neural tube duplication was not rescued in hTEM.v2 (Figure 2E).

**Figure 2.**
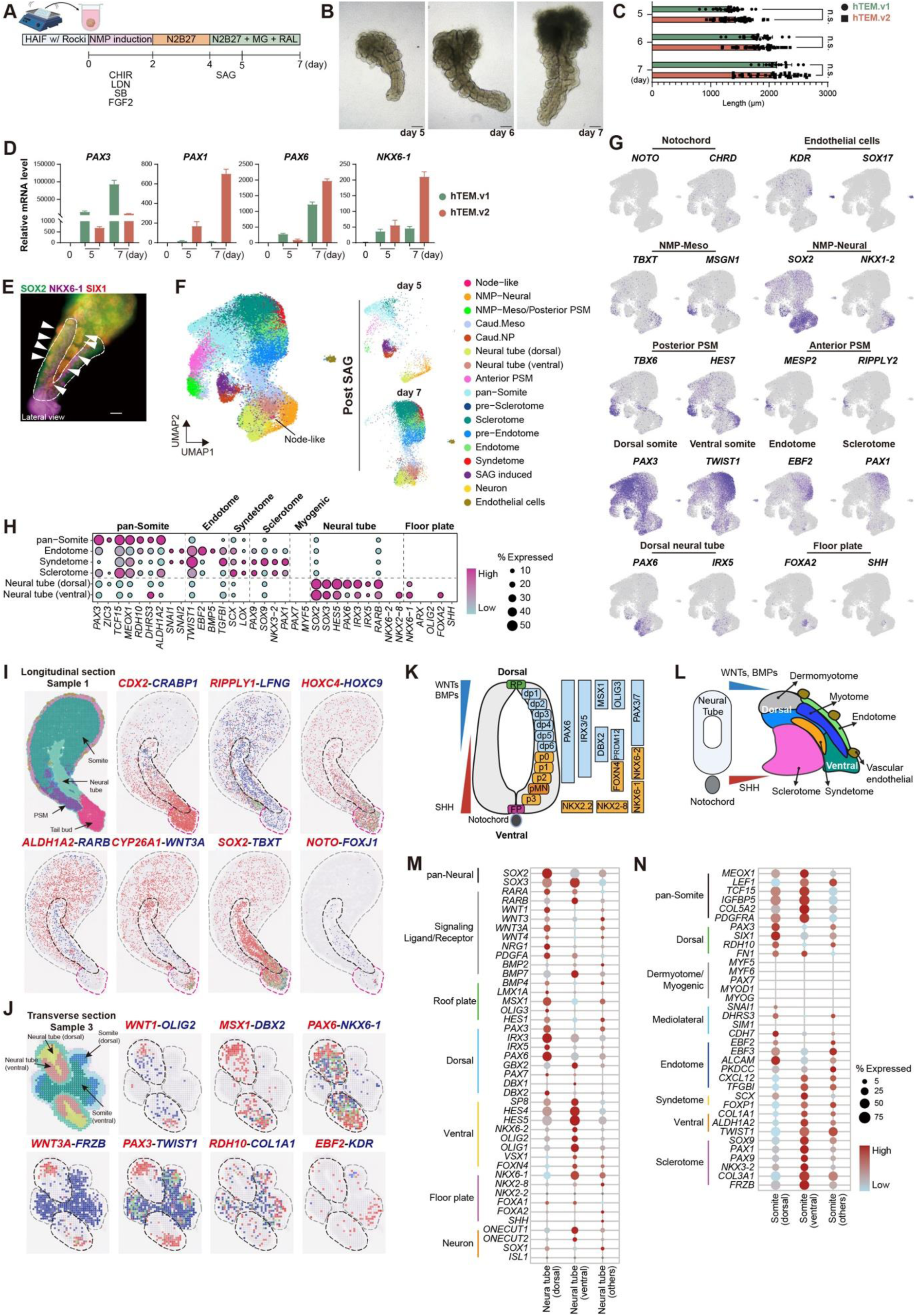
SHH activation by SAG caused D-V specifications in neural tube and somite without a notochord. (A) Schematic of hTEM.v2 generation. (B) Representative images of hTEM.v2 over days 5-7. Scale bars, 200 μm. (C) Bar plot showing length measurements comparing hTEM.v1 to hTEM.v2 over days 5-7. Each dot represents a length measurement of individual hTEM.v1/2 embryoids at shown time point (n = 14-57 for each group from same batch). n.s., no statistical significance. (D) qPCR results showing dorsal and ventral marker gene expression in hTEM.v2. Data are presented as mean ±standard deviation. Data were reproduced twice. (E) 3D projection of day-7 hTEM.v2 showing dorsal and ventral neural tubes (SOX2) flanked by paired somites (SIX1). Arrowhead, somites. Dashed line, duplicated neural tubes. Scale bar, 100 μm. (F) UMAP showing identified cell types from hTEM.v2 (days 5/7, total of 16,535 cells). Day-4 hTEM.v1 scRNA-seq data was integrated for accurate trajectory annotations. (G) UMAP showing the expression of indicated lineage markers in hTEM.v1. (H) Dot plot showing the expression profile reflecting somite compartments and neural tube D-V patterning, induced by SAG treatment. Proportions below 5% were omitted. (I-J) Spatial gene expression patterns in longitudinal (I) and transverse (J) day-7 hTEM.v2 sections. The major tissue trajectories were consistent with those shown in Figures S2E and S2G. (K-L) Schematic diagram of D-V patterning of the neural tube (K) and adjacent compartmentalization of the somite (L). (M-N) Dot plots showing the expression profiles of D-V patterns of neural tube (M) and somite (N) cells from transversely sectioned day-7 hTEM.v2 Visium HD data. The listed clusters were consistent with those in Figure S2G.

Through scRNA-seq analysis, we observed distinct ventral cell identities in hTEM.v2, including ventral neural tube (*NKX6-1*, *NKX2-8),* floor plate (*FOXA2*), ventral somite (*SNAI2*, *TWIST1*), syndetome (*SCX*), and sclerotome (*PAX1*/*9*, *SOX9*), alongside dorsal (*PAX6*) neural and dorsal (*PAX3*) somite cells (Figures 2F-2H, S2C, and S2D). However, notochord (*NOTO*, *CHRD*) cells remained limited in number (Figure 2G). This confirmed that dorsally-biased patterning in hTEM.v1 was switched to a D-V balanced patterning in hTEM.v2, due to SAG-induced SHH activity.

Attention was then turned to evaluate bi-axial patterning in hTEM.v2. Using Visium HD technology, transverse and longitudinal sections of day-7 hTEM.v2 spanning the A-P axis were acquired (Figures 2I, 2J, and S2E-S2H). To validate A-P axis formation, spatially resolved developmental events were assessed, including symmetry breaking, somite segmentation and inter-tissue RA signalling crosstalk. As seen in human embryos, the tailbud-to-hindbrain A-P patterning was evident by tailbud cells expressing *CDX2* and anterior neural cells expressing *CRABP1* (Figure 2I).^36^ The somite determination front was marked by co-expression of *RIPPLY1* and *LFNG* in the PSM region (Figure 2I).^37^ Mutually expressed *TBX18* (rostral) and *UNCX* (caudal) were observed (Figure S2I), indicative of somite segmentations.^38,39^ *ALDH1A2* (RA synthesis) expression in somite and *RARB* (RA effector) expression in neural tube cells were suggestive of somite-neural tube crosstalk along the A-P axis (Figure 2I).^40^ The anterior expression of *ALDH1A2* was opposed to posterior *CYP26A1* (RA degradation) (Figure 2I), consistent with “source and sink” RA signaling patterns that are fundamental to A-P axis establishment in chick and mouse embryos.^41,42^ As seen in previous scRNA-seq results (Figure 2G), few node-like (*NOTO*, *FOXJ1*) cells were found adjacent to bi-NMPs in the tailbud (*SOX2*, *TBXT*) (Figure 2I). Moreover, caudal expression of *HOXC9* and anterior *HOXC4* were noted (Figure 2I), indicating the presence of an emerging HOX code.^43^

Next, D-V patterning of neural tube structures in transversely sectioned hTEM.v2 was confirmed by visualizing the spatially restricted expression of roof plate (*WNT1*, *MSX1*), dorsal neural tube (*PAX6*, *DBX2*) and ventral neural tube (*OLIG2*, *NKX6-1*) markers (Figure 2J). Likewise, dorsal somite (*PAX3*, *RDH10)*^44^ and ventral somite markers (*TWIST1*, *COL1A1*) exhibited a dorsolateral-ventromedial pattern within the bilateral somites (Figure 2J). Of note, endotome (*EBF2*) and endothelial cells (*KDR*) were found in the lateral somite compartment, as observed in hTEM.v1 (Figures 2J and S1O).

These results confirmed the embryo*-*like cell-cell organization along the D-V axis for neural and somitic lineages (Figures 2K-2N). Important signals, including WNTs (*WNT1*/*3*/*3A*/*4*), BMPs (*BMP4/7*), PDGF (*PDGFA*) and heregulin (*NRG1*) displayed dorsally enriched expression patterns in hTEM.v2 (Figure 2M), similar to that observed in the neural tube *in vivo*.^45–48^ However, the floor plate (*FOXA1/2*, *SHH*) identity was under-represented (Figures 2H and 2M), suggesting an incomplete D-V axis establishment in hTEM.v2. Although *WNTs* were expressed in hTEM.v2 and are known to induce myogenesis *in vivo* and *in vitro*,^46,49^ the excessive expression of *FRZB* (water-soluble WNT antagonist) from the ventral somite cells accounts for the absence of dermomyotome and myogenic populations (Figures 2H, 2J, and 2N).

Altogether, in the absence of a notochord, exogenous SHH activation established only a limited D-V axis in hTEM.v2 and failed to rescue neural tube duplication. These observations emphasize the critical role of the notochord in trunk development and for correct D-V patterning in somitic and neural lineages.^50,51^ To establish a bi-axial embryoid comparable to the posterior trunk in human embryos, the next challenge was to establish suitable culture conditions that support co-development of notochord progenitor cells with bi-NMP descendants.

### Co-development of notochord, neural tube and bilateral somites

During gastrulation, the coordinated emergence of notochord with bi-NMPs depends on sustained WNT signaling and temporal modulation of NODAL and BMP signaling activity.^14,15,52^ Following initial WNT activation in the anterior primitive streak (APS), notochord induction coincides with the activation of a NODAL autoregulatory loop, including CER1 and/or LEFTY2.^15,53^ This precise modulation was not achieved in hTEM.v1/2, where broad and persistent NODAL inhibition by SB disrupted the APS-derived notochord process. Notably, CER1 expression precedes notochord formation and localizes to notochord adjacent APS, definitive endoderm (DE), visceral endoderm (VE) and axial progenitor populations in gastrulating mouse^54^ and human^20^ embryos (Figures S2J-S2L). Furthermore, the co-development of notochord and bi-NMPs are involved in initiation of D-V axis establishment, which depends on opposing signals of SHH and BMP2/4/7 that are also active within these early populations (Figures S2J–S2L).^55,56^

Since these critical signals were not intact in hTEM.v1 (Figure S1L), we hypothesized that its notochord deficiency resulted from; (i) disruption of APS-derived notochord induction by early and persistent NODAL inhibition (SB); (ii) absence of opposing SHH and BMP signals necessary for D-V patterning and notochord/bi-NMP specification.

To test this hypothesis, we replaced SB with recombinant human CER1 for the first 24 hours of bi-NMP induction (Figure 3A). By day 2, it was evident that temporal NODAL modulation by CER1 followed by SB supported the balanced co-emergence of APS (*OTX2*, *EOMES*), early notochord (*NOTO*, *FOXA2*), NMP-Meso (*TBXT*, *TBX6*) and NMP-Neural (*SOX2*, *NKX1-2*) progenitors (Figures S3A and S3B). Further interrogation of transcriptomic profiles in gastrulating mouse and human embryos revealed an array of signaling pathways with synergistic and opposing activity including pathways of SHH^55,56^, BMPs^47,55–58^, FGFs^12,57,59–62^, and RA^13,16,58,60^ within notochord-adjacent populations, such as APS, DE, VE and axial progenitors (Figures S2J–S2L). Through empirical testing, we established a cocktail comprised of SHH, BMP2/4/7 at days 2-4, plus temporal FGF2/3/4/8b/17 and RA at day 2-3 (Figures 3A and S3C), which supported the coordinated progress of notochord (NOTO), somitic (TBX6) and neural (NKX1-2) lineages to day 4 (Figure S3D). At day 7, markers (*SHH*, *FOXA2*) for notochord and floor plate were highly expressed in these embryoids in contrast to hTEM.v1/2 (Figure S3E). Day-7 embryoids elongated to ∼2 mm in length and formed 6–8 pairs of somites flanking a midline neural tube (Figures 3B, 3C, and S3G). We refer to these embryoids as hTEM.v3.

**Figure 3.**
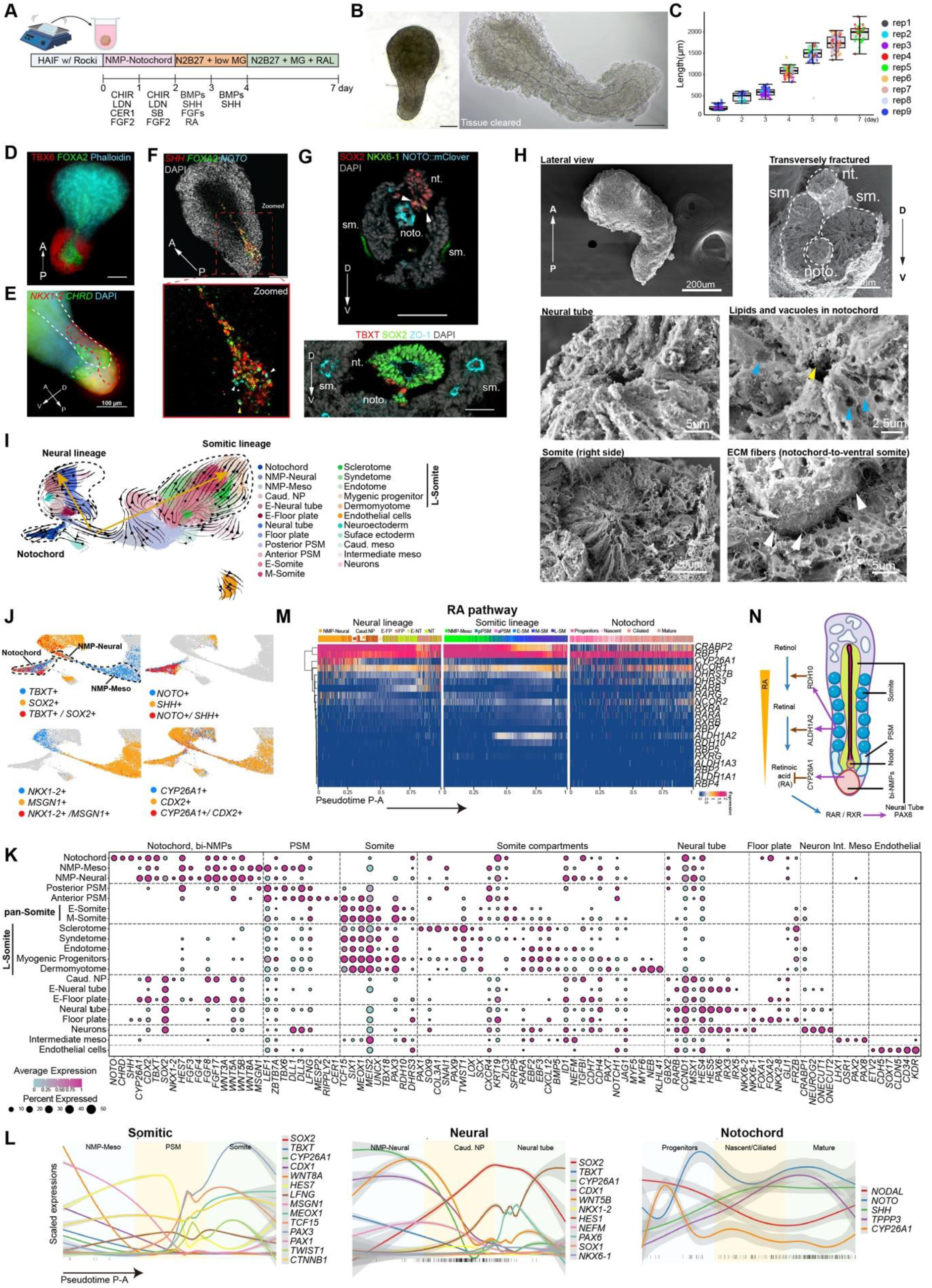
Co-development of notochord, NMP-Neural, NMP-Meso and subsequent neural and somitic lineages in hTEM.v3. (A) Schematic of hTEM.v3 generation. (B) Representative images of hTEM.v3 at day 4 (left) and day 7 (right). (C) Boxplot showing length measurements of hTEM.v3 from days 0-7 (n = 21-84 for each time point) from 9 independent biological replicates (rep). Each dot represents an individual hTEM.v3 at shown time point. (D) Immunofluorescence image showing the A-P elongating notochord (FOXA2), caudal and bilateral PSM (TBX6) and apical junctions (Phalloidin) within anterior cells. Scale bar, 100 μm. (E) HCR 3D projection showing the D-V layers of caudal neural plate cells (NKX1-2) and notochord cells (CHRD) in day-4 hTEM.v3. Scale bar, 100 μm. (F) HCR image showing transcripts of key genes (*NOTO*, *SHH*, *FOXA2*) expressed in notochord cells in day-4 hTEM.v3. The posterior node area was zoomed. Yellow arrow, triple positive. White arrow, double positive. Scale bar, 100 μm. (G) Top, Immunofluorescence images of transversely sectioned day-5.5 hTEM.v3 showing dorsally positioned neural tube (SOX2) and ventrally positioned notochord (NOTO:mClover3). Arrowhead, SOX2+NKX6-1+ ventral neural cells. Bottom, D-V patterned neural tube (SOX2) and ventral notochord (TBXT) and bilateral somites (ZO-1). nt., neural tube. sm., somite. noto., notochord. Scale bars, 100 μm. (H) SEM images of day-5.5 hTEM.v3 (n = 3). Wholemount view (top left), transversely fractured viewpoint (top-middle). Zoomed views of the transversely fractured sample showing; somite (top-right), neural tube (bottom-left), notochord with lipids (yellow arrows) and vacuoles (blue arrows) (bottom middle) and ECM fibers (white arrow, bottom-right). (I) RNA velocity analysis based on the UMAP of hTEM.v3 (days 3-7) scRNA-seq integration (total of 100,370 cells). Black arrows indicate calculated differentiation directions. Yellow arrows indicate the two major developmental streams stemming from bi-NMPs. (J) Differential expression of key genes to distinguish tailbud (*CDX2*, *CYP26A1*)-derived NMP-Neural (*SOX2*^high^T*BXT*^low^, NKX1-2), NMP-Meso (*SOX*2^low^*TBXT*^high^, *MSGN1*) and notochord (*NOTO*, *SHH*). (K) Dot plot showing the expression of identified cell type markers in hTEM.v3. Proportions below 5% were omitted. (L) Normalized expression of respective lineage marker genes along the pseudotime inferred A-P axis. Data are shown as coloured smooth spline with standard deviation in grey shade. Vertical black lines at the bottom of the plots show cells co-expressing *SOX2* and *TBXT*. (M) Heatmap showing scaled expression of genes in the RA pathway along the pseudotime inferred A-P axis. (N) Schematic diagram of RA signaling along the A-P axis of hTEM.v3.

### Morphological characteristics of hTEM.v3

Using a NOTO:mClover3 H9-hESC line, time-lapse imaging captured notochord morphogenesis in hTEM.v3 at days 3-4. This began with a salt & pepper pattern of NOTO:mClover3 expression followed by axial elongation of NOTO expressing cells, near the caudal end (Figure S3H). This aligns with *in vivo* observations that *Noto* is expressed in the node and nascent notochord in mice.^63^ Meanwhile, NKX1-2:mScarlet+ cells were progressively enriched along the midline of hTEM.v3, indicative of caudal neural plate morphogenesis (Figure S3I).

In day-4 hTEM.v3, distinct FOXA2+ notochord formed along the midline, flanked by caudal PSM (TBX6) cells (Figure 3D). Immunostaining-HCR results further confirmed the embryo-like arrangement of a dorsally-localized caudal neural plate (SOX2, *NKX1-2*) and a ventrally localized notochord (transcripts of *NOTO*, *CHRD SHH* and proteins of NOTO:mClover3, TBXT, FOXJ1, FOXA2) in hTEM.v3 (Figures 3E, 3F, S3I, and S3J).

These features closely resemble the gastrulating human CS8-9 embryos.^20,27^ After day 4, hTEM.v3 exhibited embryo-like D-V arrangements of the neural tube, notochord and bilateral somites. The emergence of a ventral notochord coincided with rescue of the neural tube duplication defect seen in hTEM.v1/2 (Figure 3G).

In mice, the node is located at the anterior tip of APS and is composed of columnar epithelial cells in the dorsal region and teardrop-shape, ciliated cells in the ventral part.^64^ SEM imaging of transversely fractured day-4 hTEM.v3 revealed a group of ciliated, squamous cells on the ventral side near the posterior end, resembling the ventral node structure in embryos (Figure S3K). Confocal imaging further confirmed ciliated cells (FOXJ1, ARL13B) residing in the presumptive node (TBXT) region (Video S2). At day 5.5, SEM of hTEM.v3 revealed a dorsally positioned lumen representing the neural tube, a vacuolated and ventrally localized notochord and bilateral rosette patterns formed by cells in the somites (Figure 3H). Consistent with observations in frog, rabbit and chick^65–67^, the inner canal of hTEM.v3 notochord contains lipid droplets of varying sizes, while the outer layer is rich in extracellular matrix (ECM) fibers, making notochord cells structurally distinguishable from neural tube cells. Notochord cells in hTEM.v3 were flattened, vacuolated and larger than neural tube cells, indicative of changes in the nature of the cytoplasm during notochord maturation. This is reminiscent of morphological features of the chicken notochord at Hamburger & Hamilton stage 14.^66^ The structural similarities between notochord structures in hTEM.v3 and frog, rabbit, chick and mouse embryos, signifies the conservation of notochord development across vertebrate species. It also validates the hTEM.v3 as a legitimate model for human trunk development.

### Cellular composition of hTEM.v3

scRNA-seq integration and clustering analysis of hTEM.v3 (days 3-7) identified 29 major cell types (Figures S3L and S3M). RNA velocity analysis further unveiled intricate developmental trajectories of the tailbud (*CDX2, CYP26A1*) stemming from notochord (*TBXT^high^SOX2^low^*, *NOTO*, *SHH*), NMP-Meso (*TBXT^high^SOX2^low^*, *MSGN1)* and NMP-Neural (*TBXT^low^SOX2*^high^, *NKX1-2*) (Figure 3I and 3J).^20,27^ Bi-NMPs diverge into two streams, including; (i) NMP-Meso → posterior PSM → anterior PSM → E-/M-Somite (pan-somite) → L-Somite (somite compartments comprised of sclerotome, syndetome, endotome, myogenic progenitors, and dermomyotome) and; (ii) NMP-Neural → Caud. NP → E-Neural tube and E-Floor plate → Neural tube and Floor plate (Figures 3I and 3K). In contrast to hTEM.v2, distinct myogenic progenitors (*PAX7*), dermomyotome (*MYF5*, *MYF6*) and floor plate (*FOXA1/2*, *NKX2-8*) cells were observed in hTEM.v3, highlighting the importance of notochord for control of cell fate specification along the D-V axis.

To better understand human notochord development, sub-clustering, RNA velocity and pseudotime analyses were conducted on the ‘notochord’ subset from hTEM.v3 (Figures S3N-S3P). Four cell subtypes delineating notochord maturation were identified (Figures S3Q and S3R). In line with APS-derived notochord processes,^15,17^ the first subtype designated as ‘node or notochord progenitors’ was marked by *NODAL*, *WNT3A*, *CYP26A1* and *CDX2*. The second and third subtypes were both marked by *NOTO*, equivalent to newly generated notochord cells *in vivo*.^63^ The second subtype was enriched for ‘nascent notochord’ (*FOXA2*, *CHRD*, *SHH*) markers,^20^ while the third subtype was signified by ‘ciliated notochord’ (*FOXJ1*, *RFX2, TCTEX1D1*) markers.^68^ In the fourth ‘mature notochord’ subtype, *NOTO* transcript was decreased associated with upregulation of *FOXA1*, *SOX9*, *NOG* and *SEMA3C*.^14^ hTEM.v3 is therefore a platform on which the detailed processes of human notochord development and function can be explored.

### hTEM.v3 recapitulates key aspects of the trunk A-P axis

To assess whether hTEM.v3 could model aspects of *in vivo* A-P axis at the multi-tissue level. Pseudotime analysis was individually performed on bi-NMP-derived somitic and neural lineages and notochord cells. We ordered cells according to the rank of pseudotime indices and inferred the A-P axis for each lineage (Figures S4A and S4B). As expected, expression of marker genes for respective lineage progenitors were enriched at the inferred posterior end, whereas marker genes for differentiated tissues were highly expressed at the anterior end (Figure 3L).

Having built the inferred A-P axis, we then assessed the spatial distribution of RA, FGF, WNT and NOTCH pathways that are crucial for trunk development as observed in vertebrate embryos.^16,42,69,70^ RA signaling is important for the balanced bi-NMP specification into somitic and neural lineages.^71^ It was therefore important to establish if RA signaling was active in hTEM.v3. The expression of RA degradation enzyme (*CYP26A1*) peaked in the posterior-most NMP-Neural and notochord progenitor cells, with RA receptor gamma (*RARG*) exclusively expressed in the NMP-Neural cells (Figure 3M). Expression of RA synthesis genes (*RDH10*, *ALDH1A2*) were restricted to anterior somitic cells and the positionally parallel expression of *RARB* was exclusive to the anterior neural cells, coinciding with the expression of *PAX6* (Figures 3L and 3M).^72^ The A-P patterned RA circuit genes confirms the integrity of RA signaling in hTEM.v3 as observed in mouse embryos (Figure 3N).^16,70^

How FGFs exert differential roles in coordinating multi-tissue co-patterning along the A-P axis remains unknown.^59,70^ Distinct expression patterns of FGFs were noted among different cell types in hTEM.v3. For example, *FGF3*/*4*/*8*/*17*/*19* in posterior neural and somitic cells, *FGF13* in anterior neural cells, *FGF13*/*18* in anterior somitic cells, while *FGF8*/*17* were expressed throughout the notochord without A-P polarity (Figure S4C).

Like FGFs, WNT ligands also showed A-P graded expression in somitic and neural lineages. Canonical (*WNT3A*/*8A*) and noncanonical (*WNT5A*/*5B*) WNT ligands were expressed in the posterior-most bi-NMPs (Figure S4D). In contrast, *WNT3A*/*5B* were expressed throughout the notochord lineage. Gradients of *CTNNB1* (WNT effector) in each lineage was opposed to the posteriorly restricted WNT ligands, consistent with polarized cell proliferation and movements during axial elongation. Anterior expression of *SFRP1*/*2* (WNT inhibitors) in both neural and somitic lineages were opposed to the WNT ligands in bi-NMPs, consistent with a negative WNT feedback loop during trunk formation in mice.^1^ How the interacting gradients of FGF and WNT signaling drive notochord formation, segmentation clock and neurogenesis in humans is not well understood hTEM.v3 will be useful for addressing such important developmental questions.

Next, we investigated the patterning of NOTCH signaling in hTEM.v3 as it is critical for axial elongation.^64,73^ How NOTCH signaling participates in D-V patterning is poorly understood, but hTEM.v3 is likely to be a useful tool to address this. In the somitic lineage, anteriorly expressed NOTCH modulator *LFNG* was opposed the posterior NOTCH ligands (*DLL3*) and its effector (*HES7*) (Figure S4E), mirroring the anterior-to-posterior somitogenesis.^42^ Although NOTCH receptors (*NOTCH1*/*2*/*3*) were lowly expressed in the neural and notochord lineages, NOTCH effectors (*HES1*/*4*) and their target (*CCND1*) were highly expressed in anterior neural cells and throughout the notochord, respectively. This could explain the rapid morphogenesis and axial elongation of neural tube and notochord after day 4 (Figures 3B and 3C).^74,75^ Interestingly, we noted the expression of a NOTCH coactivator *MAML2* throughout the notochord (Figure S4E). The role of *Maml2* is possibly involved in Sox9-dependent inhibition of the WNT pathway in mouse sclerotome.^76,77^ hTEM.v3 offers a unique opportunity to characterize this notochord-dependent NOTCH regulatory mechanism in human ventral patterning.

Collectively, hTEM.v3 faithfully recapitulates A-P axis development at multi-tissue levels, mirroring key signaling networks of early human embryogenesis. By achieving high-fidelity reconstruction of essential signaling pathways, this model opens new avenues to interrogate spatiotemporal signaling crosstalk and cell-fate decisions at a multi-tissue level.

### Patterning of neural and somitic cells along the D-V axis

To delineate D-V specifications in hTEM.v3, sub-clustering and RNA velocity analysis was performed on neural and somitic cells, respectively (Figures 4A and 4B). The resultant UMAP demonstrated clear D-V patterning, evident by distinct transcriptomic profiles of dorsal/ventral neural cells-floor plate and somite compartments, respectively (Figures 4C, 4D, S5A, and S5B). This high degree of complexity regarding D-V specifications was not seen in hTEM.v1/2 (Figure 4E). Immunostaining of transversely sectioned day-6.5 hTEM.v3 confirmed D-V patterned dorsal (PAX6) and ventral (NKX6-1, OLIG2) neural cells along the neural tube (Figure 4F). These midline-positioned neural cells were flanked by distinct somite compartments including the dorsal somite (PAX3), dorsolateral dermomyotome (MYF5:mClover3), myogenic progenitors (PAX7), lateral endotome (EBF2:mScarlet), ventromedial sclerotome (PAX1) and surrounding vascular endothelial cells (SOX17) (Figures 4F and 4G).

**Figure 4.**
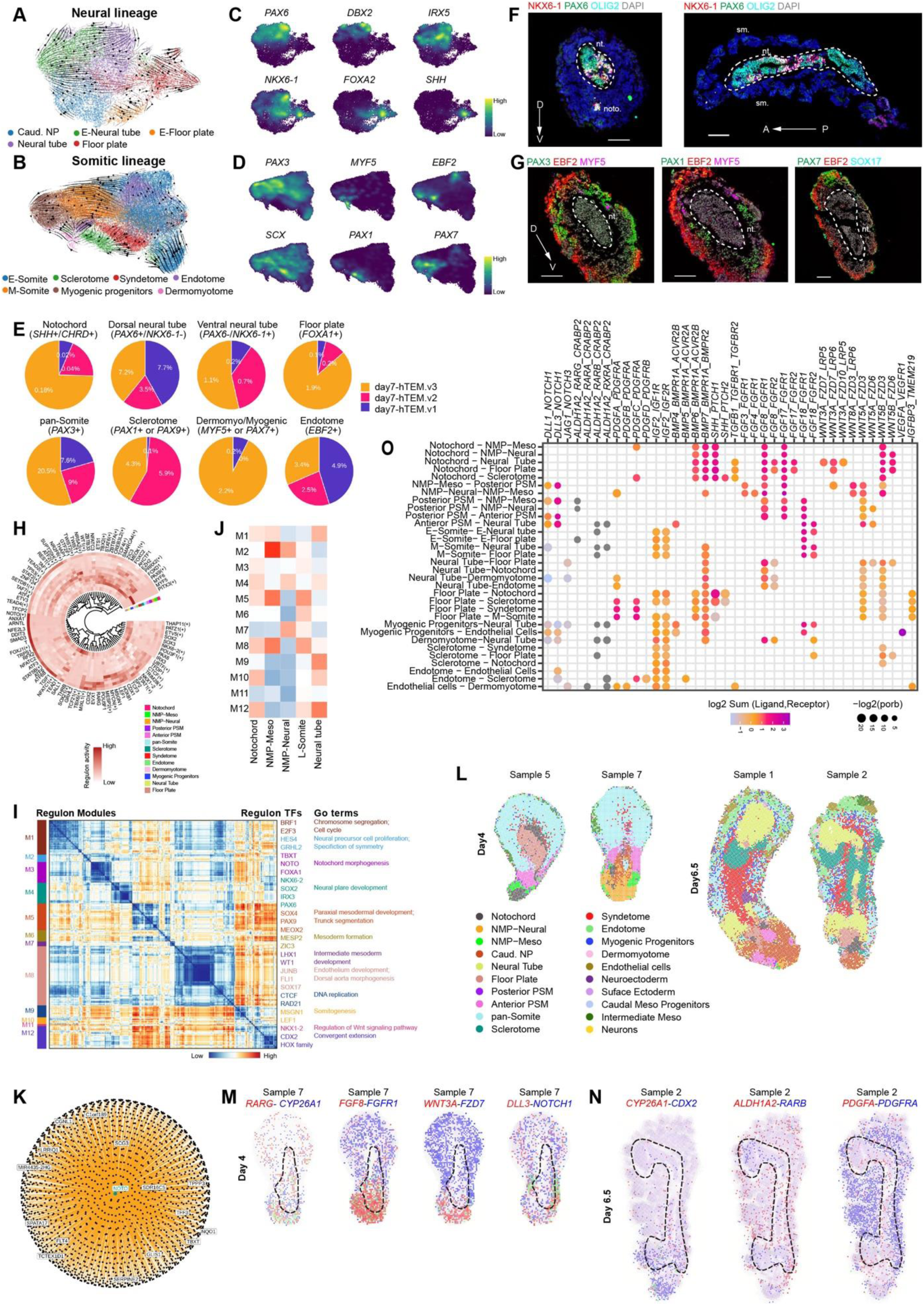
Molecular profiling of D-V organization and spatial patterning for hTEM.v3. (A-B) RNA velocity analysis of neural (A) and somitic (B) lineage subclusters from hTEM.v3 (day 3-7) scRNA-seq integration in Figure S3L. To distinguish somite compartments, regression was performed on TBX18 and UNCX to remove somite A-P patterns before re-clustering. (C-D) Density plot showing trajectories of D-V patterned neural (C) and somitic (D) lineages in hTEM.v3. Dorsal neural markers; *PAX6*, *DBX2*, and *IRX5*. Ventral neural and floor plate markers; *NKX6-1*, *FOXA2* and *SHH*. pan- and dorsal somite; *PAX3*. Dermomyotome; *MYF5*. Endotome; *EBF2*. Syndetome; *SCX*. Sclerotome; *PAX1*. Myogenic progenitors; *PAX7*. (E) Comparison of hTEM.v1/2/3 cell types to contrast the presence or absence of D-V fates. (F) Immunofluorescence images of transverse (left) and longitudinal (right) sections of day-6.5 hTEM.v3 showing the presence of D-V patterning in the neural tube structure along D-V and A-P axes. Dorsal neural marker; PAX6. Ventral neural and floor plate markers; OLIG2 and NKX6-1. Scale bars, 100 μm. (G) Immunofluorescence images of transverse sections of day-6.5 hTEM.v3, showing somite compartments along D-V axis. pan- and dorsal somite, PAX3. Endotome, EBF2:mScarlet. Dermomyotome, MYF5:mClover3. Myogenic progenitors, PAX7. Endothelial cells, SOX17. Scale bars, 100 μm. (H) Circular heatmap showing the regulon representing TF genes in the indicated cell types from hTEM.v3. (I) Heatmap (left) of regulon modules based on cell type associated regulon activities in Figure S5C, with the corresponding GO terms related to each regulon module highlighted (right). (J) Heatmap showing the enrichment of regulon modules across listed cell types. L-Somite, late somitie, including sclerotome, syndetome, dermomyotome, myogenic progenitors and endotome cells from hTEM.v3 scRNA-seq data in Figure S3L. (K) *NOTO* gene regulatory networks in hTEM.v3. The top 15 interacting genes are displayed. (L) Spatial UMAP of RCTD cell type annotations in day-4 and day-6.5 hTEM.v3 (n = 8 each). Colors for annotation and cell types displayed are consistent with those in Figure S6B-S6D. (M-N) Spatial gene expression patterns in longitudinally sectioned day-4 (M) and day-6.5 (N) hTEM.v3. Dashed lines, areas for neural plate (day 4) and neural tube (day 6.5). (O) Bubble plot showing the selected ligand-receptor interactions between indicated cell types from hTEM.v3. Dot color represents the normalized sum of expression level of ligand in source cells and interacting receptor in targeting cells.

To understand the molecular mechanisms underlying D-V axis establishment, we performed Gene Ontology (GO) and pathway enrichment analysis on scRNA-seq data for hTEM.v3. First, SCENIC^78^ was used to generate a regulon module enrichment heatmap illustrating representative transcription factor (TF) genes associated with all cell types of hTEM.v3 (Figures 4H, 4I, and S5C). As expected, biological processes in GO terms enriched for each regulon module were consistent with associated cell types (Figure S3L, S5D, and S5E). For example, module M4 comprised of key TFs (*PAX6*, *IRX3*, *NKX6-2*) important for regulating neural tube formation, was significantly enriched in “GO:0001840 neural plate development”, was highly expressed in the “Neural tube”, but not in “NMP-Meso” or “L-Somite” clusters in hTEM.v3 scRNA-seq data (Figures 4I and 4J).

Regulon module-based gene regulatory network (GRN) analysis identified key GRNs (*NOTO*, *PAX6*, *NKX6-2*, *FOXA1*/2) and their target networks that drive D-V axis formation (Figures 4K and S5F). The *NOTO* GRN was enriched for SHH signaling (*GLIS1*), *TBXT,* and cilia functions (*TPPP3*, *SCG3*, *TCTEX1D1*). Consistent with Noto’s role in mice,^79^ *NOTO* target genes were expressed in human ’nascent’ and ’ciliated’ notochord subtypes (Figure S3Q). The NKX6-2 GRN contained targets that balance ventral neural tube patterning, including interneuron (*HES5*)^80^ and motoneuron specifiers (*OLIG1*/*2*)^81^ (Figure S5F). Finally, *FOXA1*/*2* GRNs exhibited differential functions. *FOXA1* targets were associated with floor plate (*ARX)*^82^ patterning and notochordal fluid trafficking (*CFTR)*^83^, whereas the *FOXA2* targets governed node formation (*PPIL6*)^84^, cilia function (*CFAP43*)^85^, and maintaining notochord structure (*KRT8*, *EPCAM*, *FN1*)^63^ (Figure S5F). The GRN analysis revealed multifaceted networks in ventral patterning, demonstrating the establishment of D-V axis in hTEM.v3.

Overall, hTEM.v3 self-organizes into notochord, neural tube and somitic tissues with proper A-P and D-V patternings (Figures 3E-3H and 4A-4G). A key advance in hTEM.v3 is its ventral neural specification. Unlike hTEM.v2, which displayed elevated *HES5* and low *NKX6-2* expression accompanied with a floor plate deficiency (Figure 2H), hTEM.v3 exhibited distinct floor plate (ARX) and ventral neural (OLIG2) markers alongside reduction in *HES5* and elevated *NKX6-2* (Figure 3K). Our GRN analysis suggests that NXK6-2 acts as a pivotal node, connecting SHH signaling and Notch-mediated HES5 activity to balance motoneuron (*OLIG2*)/interneuron (*HES5*) specification (Figures S5A and S5F).^86,87^ These data suggest a previously undescribed SHH-NKX6-2-HES5 regulatory mechanism in human neural tube patterning.

### Spatially transcriptomic profile reveals embryo-like signatures of hTEM.v3

To map spatial allocation of cells in hTEM.v3, we performed Visium HD on longitudinal sections (days 4/6.5). Key cell types including notochord, NMP-Meso, NMP-Neural and D-V fated neural and somitic cells were all identified (Figure S6A). However, full recovery of D-V fated cells from Visium HD data, especially in somite compartments, was difficult due to limited sections. We therefore performed cell type deconvolution (RCTD)^88^ by leveraging the reference annotations from scRNA-seq data of hTEM.v3 (days 3-7) (Figure S6B). The resulting spatial UMAP exhibited embryo-like spatial patterns that correspond to anatomical regions of the posterior trunk in human CS8-10 embryos (Figures 4L, S6C, and S6D).^20,27,36^ For example, by day 4, the Caud. NP (*NKX1-2*, *GBX2*) extending anteriorly out of the tailbud was flanked by anterior PSM (*RIPPLY2*, *CER1)*^89^ (Figure S6E). By day 6.5, neural tube and floor plate were specified in the midline along the A-P axis and flanked by compartmentalized somites (Figures 4L and S6D). Remarkably, NMP-Neural and NMP-Meso cell populations occupied distinct regions in the day-4 tailbud (Figure 4L, S6C and S6E), representing an unprecedented bi-layered structure of bi-NMPs *in vitro*. This bi-layered structure signifies the onset of D-V patterning as observed in a spatially resolved human CS9 embryo.^27^

Next, we verified the inferred A-P signaling gradients from spatially resolved ligand-receptor gene expression. For example, the RA (*RARG*-*CYP26A1*), FGF (*FGF8*-*FGFR1*), WNT (*WNT3A*-*FZD7*) and NOTCH (*DLL3*-*NOTCH1*) ligand-receptor pairs were spatially enriched in the tailbud of hTEM.v3 (Figures 4L and 4M), consistent with their inferred A-P distribution in Figures 3M and S4C-S4E. In addition, CellChat^90^ predicted tissue-tissue communications from scRNA-seq data that were also validated by Visium HD data (Figure 4O). For example, *PAX1* (target of SHH pathway) expression in the sclerotome region was shown to be in close proximity to *SHH* expression in the floor plate (Figure S6F), reflecting the SHH signaling range. The anterior expression of *ALDH1A2* in somites and posterior *CYP26A1* in tailbud (*CDX2*) were evident of the RA signalling along A-P axis (Figures 3M and 4N). The detection of *RARB* and *CRABP1* (RA target) in anterior neural cells confirms the somite-to-neural RA crosstalk has been established as in the spinal cord of human CS10 embryos (Figure S6F).^36^ Moreover, CellChat-predicted PDGF (*PDGFA*-*PDGFRA*) signaling between neural and somitic cells was revealed on the spatial UMAP (Figures 4N and 4O). This suggests a neural-to-somite regulatory mechanism in patterning the sclerotome.^91^ These spatial data demonstrate hTEM.v3’s utility in studying advanced organogenesis co-patterning.

The emergent HOX expression is key to axial elongation and spatially aligned with body axes formations,^92^ we therefore evaluated the fidelity of HOX coding in hTEM.v3. hTEM.v3 exhibited HOX collinearity in somitic lineage as observed in axialoids^8^ and extended it to neural and notochord lineages, with over 50% of HOX genes displaying coordinated expression (Figure S4F). Tissue-specific expression patterns of HOX genes were also noted. For example, *HOXA5* and *HOXB5* were expressed in anterior somitic and notochord lineages but not in the neural lineage. *HOXA9* was restricted to the Caud. NP and posterior PSM, but not expressed in notochord cells (Figure S4F), consistent with its initial expression in caudal neural plate region in mouse gastrulating embryos.^93^ We next exemplified the spatiotemporal *HOX* coding with scaled expression of *HOXC genes* from days 4-6.5 (Figures S6G and S6H). During axial elongation, a clear anterior expansion of *HOXC6* and *HOXC8* expression accompanied the caudal downregulation of *HOXC10,* as observed in the human developing spinal cord.^94^ Collectively, our spatial and scRNA-seq transcriptomic data establishes hTEM.v3 as a high-fidelity model of human posterior axial development. This system recapitulates the spatially patterned signaling pathways and tissue-tissue crosstalk of human posterior trunk. Importantly, hTEM.v3 demonstrated a spatiotemporal HOX coding alongside axial elongation, thereby capturing the core signaling and transcriptional machinery for future developmental studies.

### Developmental staging of hTEM.v3 resembles primate CS8-10 embryos

Next, we sought to allocate the developmental lineages of hTEM.v3 to those involved in trunk formation in primates and mice. First, a embryogenesis scRNA-seq reference, consist of public datasets of human (CS7/8/10)^20,27,84^ and cynomolgus monkey (CS8/9/11)^19^ embryos, was created based on primate orthologues (Figure S7A). Consist with hTEM.v3, the notochord, NMP-Neural and NMP-Meso in human-monkey (H-M) reference dataset showed well-matched expression profiles (Figure 3K and 5A). Next, sub-clustering of caudal trajectories (caudal epiblast, caudal mesoderm, somitic mesoderm and NMP) in mouse embryo (E7.0-E8.5) datasets^19,54^ identified previously unresolved mouse NMP-Meso and NMP-Neural populations (Figure 5B), highlighting the evolutionary conservation in trunk formation between mice and primates.

**Figure 5.**
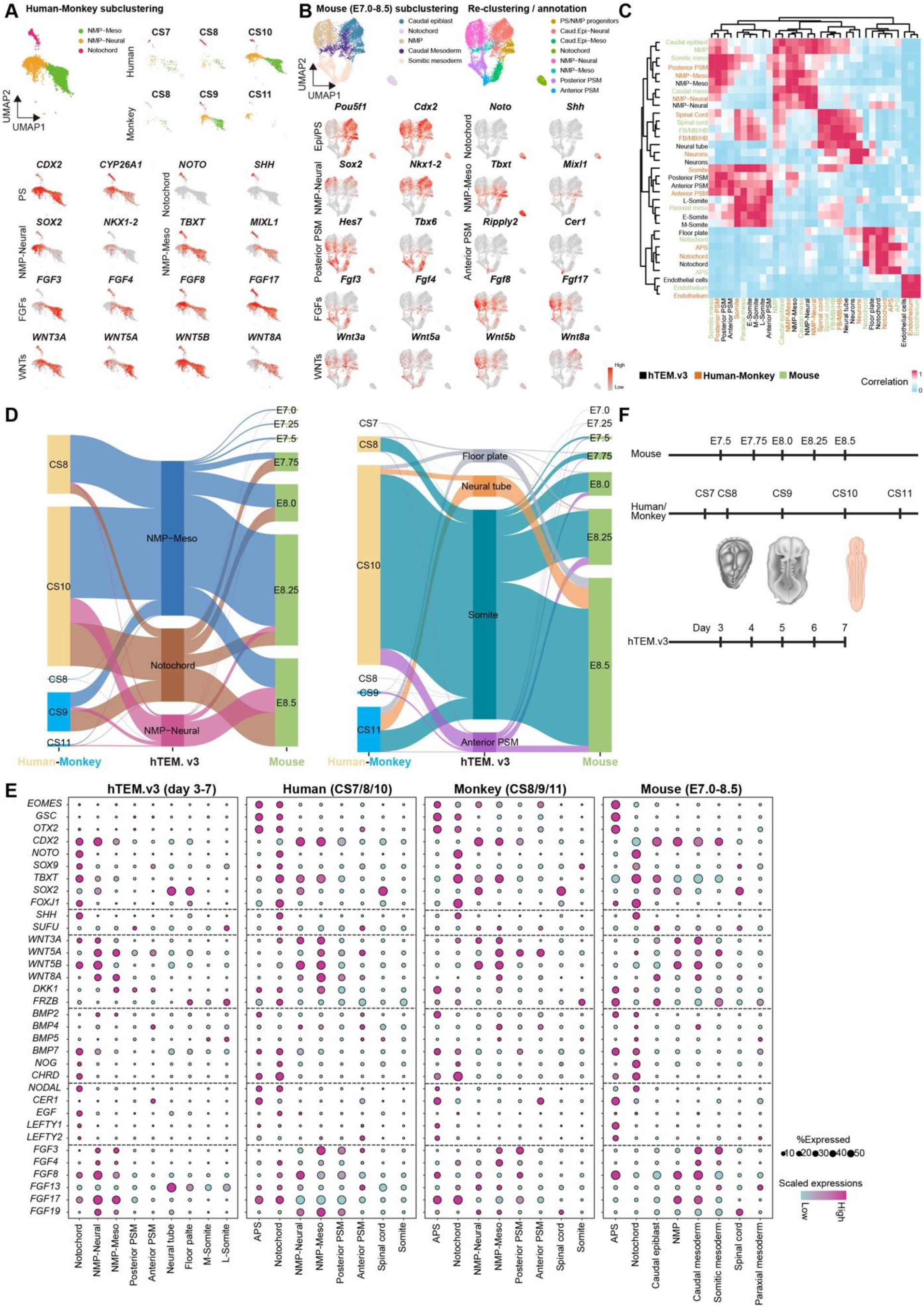
Developmental staging of hTEM.v3 resembles primate CS8-10 embryos. (A) Top, sub-clustering UMAP revealing the notochord, NMP-Neural and NMP-Meso populations from human (CS7/8/10) and monkey (CS8/9/11) embryo integration dataset. Bottom, differential gene expression patterns delineating these cell types. (B) Sub-(top-right) and re-(top-right) clustering UMAPs revealing the notochord, NMP-Neural and NMP-Meso from mouse embryo dataset (E7.0 to E8.5). Bottom, differential gene expression patterns delineating these cell types. (C) Heatmap of correlation co-efficient among cell types from human, monkey, mouse and hTEM.v3. The correlation was calculated and summarized from Figure S7F. (D) Sankey diagram showing relationships between selected cell types in hTEM.v3, human (CS7/8/10), monkey (CS8/9/11) and mouse embryos (E7.0 to E8.5). (E) Gene expression profiles for key transcription factors and key signaling pathways (SHH, WNT, BMP, NODAL, and FGF), in cells forming the posterior trunk in hTEM.v3, human, monkey and mouse embryo datasets. (F) Cartoon summary of similarities across human, monkey, mouse embryos with hTEM.v3 in developmental stages.

Next, we examined cellular composition similarities between hTEM.v3 (days 3-7) and H-M (CS7-11) datasets using Seurat. Comparative projection of cell types showed that hTEM.v3 accurately recapitulates H-M posterior trunk development including the notochord, bi-NMPs, somite and spinal cord (Figures S7B and S7C). Likewise, hTEM.v3 was mapped to ortholog mouse trajectories of axial progenitors (notochord, caudal epiblast, caudal mesoderm, somitic mesoderm, NMP), paraxial mesoderm and spinal cord (Figures S7D and S7E). Collectively, an unsupervised cluster similarity analysis summarized the major trunk cell types conserved across hTEM.v3, H-M, and mouse embryos (Figures 5C and S7F). Further comparative analysis aligned the developmental stages between hTEM.v3 and H-M, and hTEM.v3 and mouse, respectively (Figures 5D and S7G). We noted the correspondences of advanced cell types, such as neurons, intermediate mesoderm and endothelial cells at post-gastrulation stages between datasets (Figure S7G). Moreover, genes important for notochord, bi-NMP specification and pathways for RA, FGF and WNT signaling exhibited broad similarities across species (Figures 5E and S7H). One exception is the absence of *Fgf17* expression in mouse notochord. In hTEM.v3 and H-M notochord populations, FGF17 expression is highly expressed, suggesting a divergence in mechanisms of notochord progress and D-V establishment between mice and primates (Figures 5A and 5E). These findings establish that hTEM.v3 is suitable for identifying new mechanisms of human development.

In summary, combined with the embryo-like morphological features of hTEM.v3 (Figures 3D and S3I), cross-species comparative analyses indicates that hTEM.v3 accurately models cellular and molecular aspects of trunk development in CS8-10 H-M embryos. hTEM.v3 (days 3-7) equates approximately to E19-23 in humans, E20-24 in cynomolgus monkeys and E7.5-8.5 in mice (Figure 5F).

### Changes in notochord identity and SHH signaling switch D-V fates

Next, the utility of hTEM.v3 as a testbed for investigating human D-V patterning at the genetic level was explored. Focus was directed towards NOTO which is important for notochord function in mice by serving as a key regulator of Shh signaling in mice.^20,63^ Loss-of-function mutation of NOTO (NOTO-LOF) derived day-6 hTEM.v3 exhibited irregular posterior morphology and shorter axial length (Figure S8A-S8C), consistent with the shortened tail phenotype in *Noto*-null mice.^79^ Comparative scRNA-seq analysis between NOTO-LOF and wild-type (WT) revealed an expanded population of ‘mature notochord’ (*FOXA1*, *NOG*) cells in day-6 hTEM.v3 (Figures 6A–6C). This shift in notochord identity correlated with a pronounced upregulation of SHH expression and ventrally-biased fates (Figures 6C-6E). For example, cellular proportion of ventral lineages (floor plate; *FOXA2*, *NKX6-1*. sclerotome; *PAX1*, *PAX9*) were upregulated at the expense of dorsal identities in neural tube (*PAX6*, *MSX1*) and somites (*PAX3*) (Figures 6D, 6E, and S8D). Focusing on the neural tube, elevated SHH activity (*SHH*, *GLI1*) in NOTO-LOF sharply suppressed PAX6 while inducing NKX6-1 (Figures 6E and 6F). Moreover, NOTO-LOF downregulated cilia-related genes (*FOXJ1*, *RFX3*, *TCTEX1D1*) (Figure S8E), consistent with Noto’s conserved role in regulating ciliogenesis.^79^ These data demonstrate hTEM.v3 as a valuable testbed for genetic dissection of human development and establish the role of NOTO in restraining SHH activity and safeguard nascent notochord identities.

**Figure 6.**
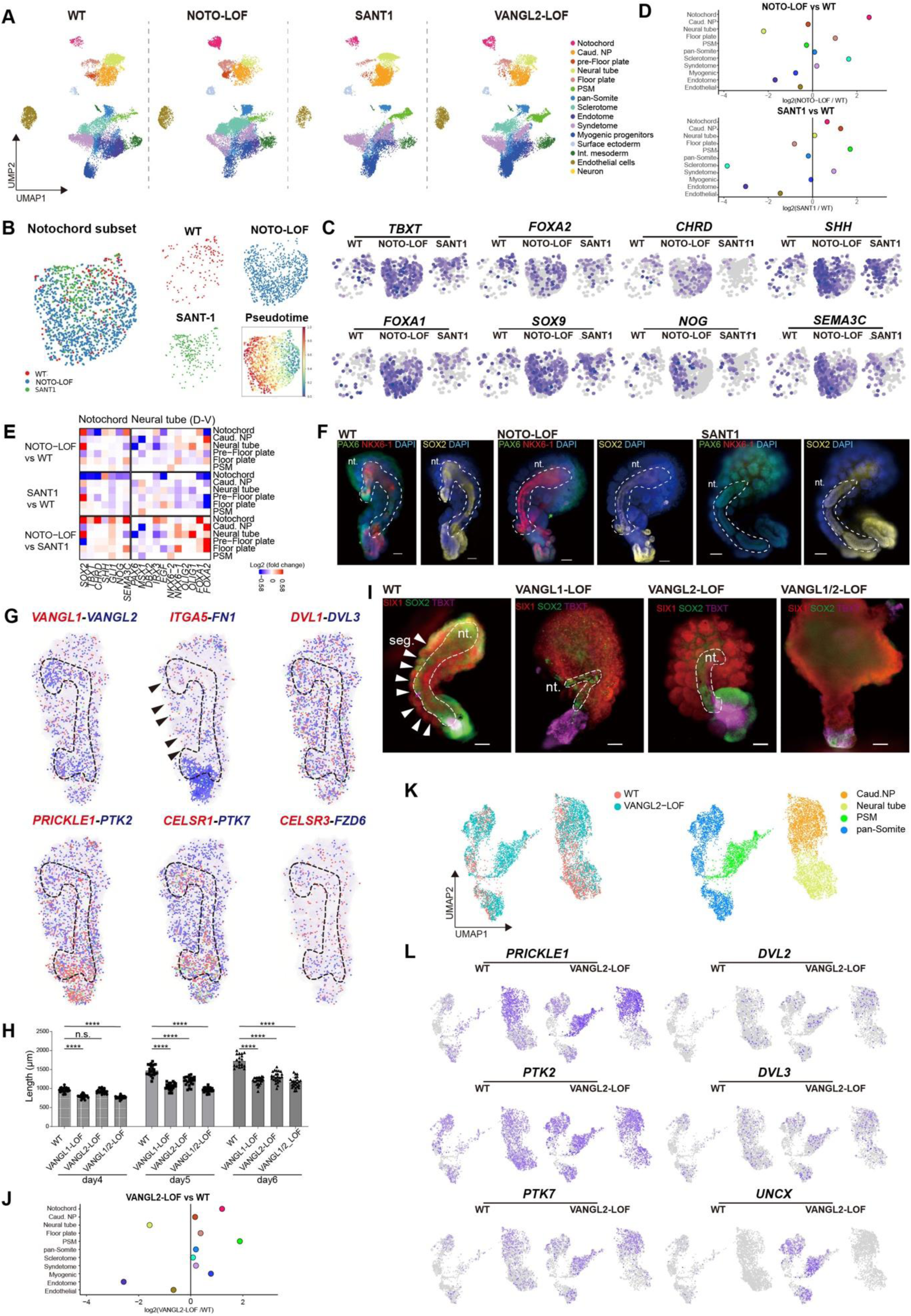
SHH signaling modulation and disease modeling using hTEM.v3. (A) UMAP of integration of respective day-6 hTEM.v3 scRNA-seq data from WT (14,462 cells), NOTO-LOF (16,081 cells), SANT1 (250 nM, days 4-6, 14,502 cells), and VANGL2-LOF (16,504 cells). (B) UMAP showing the re-analysis of notochord cells from WT, NOTO-LOF, and SANT1 in (A). Pseudotime scores reflect the developmental direction indicating notochord maturation. (B) UMAP showing expression of marker genes indicating notochord maturation. (C) Changes in cellular composition from NOTO-LOF (top) and SANT1 treatment (bottom) in contrast to WT. Dots represents log_2_(NOTO-LOF/WT or SANT1/WT) fold changes for listed cell types. (D) Heatmap showing changes in the expression of key genes in regulation of notochord development, neural tube D-V patterning, FGF and NOTCH signaling upon NOTO-LOF and SANT1 treatment. (E) Whole-mount immunostaining of dorsal (PAX6) and ventral (NKX6-1) fate changes in neural tube (SOX2) in WT, NOTO-LOF and SANT1 treated day-6 hTEM.v3. nt., neural tube. Scale bars, 100 μm. (F) Spatial expression patterns of representative PCP genes in day-6.5 hTEM.v3 (sample 2 in Figure 4L). Dashed lines, areas of neural tube. Black arrows, somite segments. (G) Comparison of length measurements using hTEM.v3 between WT, VANGL1-LOF, VANGL2-LOF and VANGL1/2-LOF (n ≥ 21 in each group at different times). ****p values < 0.0001 were calculated using students’ t-test. n.s., no statistical significance. Results were reproduced in more than 3 biological replicates. (H) Whole-mount Immunostaining of day-6 hTEM.v3 showing defects in neural tube elongation (SOX2) and somite (SIX1) segmentation using WT, VANGL1-LOF, VANGL2-LOF and VANGL1/2-LOF lines. Arrow heads, somite segment. nt., neural tube. Scale bars, 100 μm. (I) Changes in cellular composition in VANGL2-LOF vs WT. Dots represents log_2_(VANGL2-LOF/WT) fold changes for listed cell types. (J) Sub-clustering of PSM, pan-Somite, Caud.NP and Neural tube from VANGL2-LOF and WT samples shown in (A). (K) UMAP showing PCP gene expressions involved VANGL2-related human neural tube defects.

In contrast to the NOTO-LOF, exposure to SANT1^95^ (smoothened antagonist, 250 nM at days 4-6) led to a downregulation of ventral (*PAX1*, *NKX6-1*, *FOXA2*) markers without negatively impacting dorsal (*PAX3*, *PAX6*) markers or axial elongation (Figures S8B-S8D). scRNA-seq and immunostaining results confirmed that SANT1 treated hTEM.v3 were dorsally-biased similar to hTEM.v1 (Figures 6E and 6F). This was supported by reduced ventral populations (e.g., floor plate, sclerotome) and a concordant reduction in the expression of ventral markers (*NKX6-1/6-2*, *OLIG1/2*, *FOXA1/2*) (Figures 6D, 6E, and S8D). It is noteworthy that the proportion of notochord cells and endogenous levels of SHH were not negatively affected in SANT1 (Figures 6C and 6D), indicating that the dorsally-biased fates in SANT1 can be attributed to the blockade of SHH signaling between notochord and its targets. Pharmacological inhibition of the SHH pathway downregulates BMPs, VEGFs and KDR which control dorsal aorta development and vasculogenesis.^96^ In support of this observation, reductions in the proportions of endotome, endothelial cells and related gene expression (*MEF2C*, *CD34*, *KDR*) are noted in hTLE.v3 (Figures 6B, S8F, and S8G).

Collectively, these observations show that by modulating notochord identity and derived SHH signaling, hTEM.v3 embryoids can be patterned into ventrally-biased (NOTO-LOF) or dorsally-biased (SANT1) fates. This responsiveness allows timed perturbations in SHH to be assessed and enables human bi-axial development to be explored with temporal precision and quantifiable measurements.

### Disease modeling of axial truncation by knockout of *VANGL1/2*

We next evaluated hTEM.v3 as a platform for the study of human congenital disease. For proof-of-concept, PCP signaling core members VANGL1 and VANGL2 were selected, as mutations in *VANGL1*/*2* lead to NTDs including spina bifida and congenital vertebral malformations.^97–99^ As expected, *VANGL1*/*2* expression was widespread throughout day-6.5 hTEM.v3, with *VANGL2* enriched in neural tube and *VANGL1*/*2* slightly enriched in caudal neural plate and PSM (Figures 6G and S8H). To model VANGL-related defects in these tissues, loss-of-function mutations of *VANGL1*/*2* H9-hESC lines were made, including *VANGL1* ^-/-^ (VANGL1-LOF), *VANGL2* ^-/-^ (VANGL2-LOF), and *VANGL1* ^+/-^; *VANGL2* ^-/-^ (VANGL1/2-LOF) (Figure S8A).

As in mice,^99,100^ *VANGL*-mutant derived hTEM.v3 exhibited axial truncations with different degrees of penetrance (Figures 6H and S8I). VANGL1/2-LOF displayed an early failure in mediolateral narrowing on day 4 and subsequently, severe dysregulation of somitogenesis (no segmentation) and neural tube genesis (loss of a neural tube) (Figure 6I). These observations are consistent with the well-established role of *Vangl* in C&E movements and PCP signaling during axial elongation in zebrafish and mouse models.^99,100^ Contrarily, VANGL1-LOF and VANGL2-LOF showed milder axial truncation defects compared to VANGL1/2-LOF, with VANGL2-LOF reproducing the no-segmentation defect presented in VANGL1/2-LOF (Figure 6I).

To further characterize the role of VANGL2 in trunk development, we performed a comparative scRNA-seq analysis of VANGL2-LOF and WT embryoids (Figure 6A). In VANGL2-LOF, a sharp decrease in the proportion of neural tube cells was accompanied by a marked increase in PSM, while the somite population remained unchanged (Figure 6J). The expanded PSM population and the caudal accumulation of TBXT protein in VANGL2-LOF mirrored the observations of widened PSM and caudally restricted expression of *Tbxt* and *Tbx6* in *Vangl2* mutant mice (Figure 6I).^99^ Focusing on the loss of somite segmentations in VANGL2-LOF, differential gene expression analysis revealed that *ITGA5*, a gene required for fibronectin (*FN1*) accumulation during somite boundary formation and neural tube closure (Figure 6G),^101,102^ was downregulated in pan-somite cells (Figure S8J). This indicates the importance of stable interactions between VANGL2 and integrins in somite segmentation and points towards an unknown VANGL2-dependent mechanism that regulates *ITGA5* expression.^103^

Turning to VANGL2-related PCP components, *PRICKLE1, DVL1*/*3*, *PTK7*, *CELSR1*/*3* and *FZD6* were broadly expressed throughout the somite and neural tube trajectories in day-6.5 hTEM.v3 (Figure 6G). Although VANGL2 protein is typically negatively correlated with PRICKLE1/2 protein levels,^104^ VANGL2-LOF resulted in increased *PRICKLE1*/*2* transcript levels in PSM in VANGL2-LOF (Figures 6K, 6L and S8J). This finding points towards the existence of compensatory transcriptional mechanism within the PCP network in PSM cells. Additional PCP genes including *CELSR3*, *DVL2/3, PTK7* were concordantly upregulated with over-expanded *UNCX* (caudal polarity) expression in PSM and pan-somite cell populations (Figures 6L and S8J). In contrast, VANGL2-LOF caused downregulation of *PRICKLE2*, *CELSR3, DVL2* and *PTK2* in neural cells, accompanied by a shortened neural tube (Figures 6I and S8J), consistent with their reported implication in human NTDs.^105^ This contrasting expression pattern in VANGL2-LOF indicates a differential role of VANGL2 in coordinating PCP components in; (i) establishing A-P polarity somite segmentations and, (ii) neural tube elongation.

Current *VANGL*-related early congenital disease models are based solely on observations made in zebrafish and mice.^99,106^ Using *VANGL1*/*2* knockouts, we demonstrated the potential for our multi-tissue, bi-axial embryoids to serve as a NTD model and as a platform for developing genetic screens, therapeutic strategies, toxicity and drug tests in a more human relevant context.

Overall, by leveraging the co-development of notochord and bi-NMPs, embryo-like structures were generated that, at the morphologic and molecular level, recapitulating A-P and D-V axes formations in the human posterior trunk. These embryoids are self-organized and non-integrated. This provides a valuable tool that meets the criteria^2^ for benchmarking early human embryogenesis and the development of human disease models. This can also be a platform for future development of next-generation embryoids that recapitulate additional aspects of human development.

### Limitation of this study

Few to no neural crest (*SOX10*), endoderm (*HNF4A*), cardiac mesoderm (*HAND1*) or anterior neural ectoderm (*OTX2*) cells were detected in hTEMs. The neural tube structures in embryoids are believed to represent secondary neurulation, which accounts for the caudal portion of the A-P axis.^107^ More advanced embryo models are needed to reconstitute the fully body axes of A-P, D-V and L-R.

### Resource availability

The raw and processed sequencing data in this study are available in the GEO repository GSE314260. Source code for analyzing the sequencing data is available at https://github.com/tianmingwucuhk/hTEM.

## Supporting information

Key Resources Table

## Acknowledgement

We thank the Core Laboratories of School of Biomedical Sciences at the Chinese University of Hong Kong for the service of single cell sorting, histology sections, confocal imaging and SEM. This study was supported by grants to S.D. from the Research Grants Council of Hong Kong (General Research Fund) and the Hong Kong Jockey Club Charities Trust. S.D. is a Global Stem Scholar and Director of the JC STEM Lab of Stem Cells and Regenerative Medicine.

## Author contributions

Conceptualization, T.-M.W. and S.D.; Sequencing data generation, T.-M.W, H.Y., and J.V.; Sequencing data analysis, T.-M.W, H.Y., B.S.-H.W., L.X., and S.K.-W.T.; Experiments, T.-M.W., K.-X.T., W.-M.X., E.S.-K.N., A.Y.-F.K., and Jianan Z.; Interpretation and discussion, T.-M.W., S.D., Jiannan Z., B. G.; Supervision, T.-M.W. and S.D.; Manuscript writing, T.-M.W. and S.D.; Funding acquisition, S.D.

---

**Figure S1.**
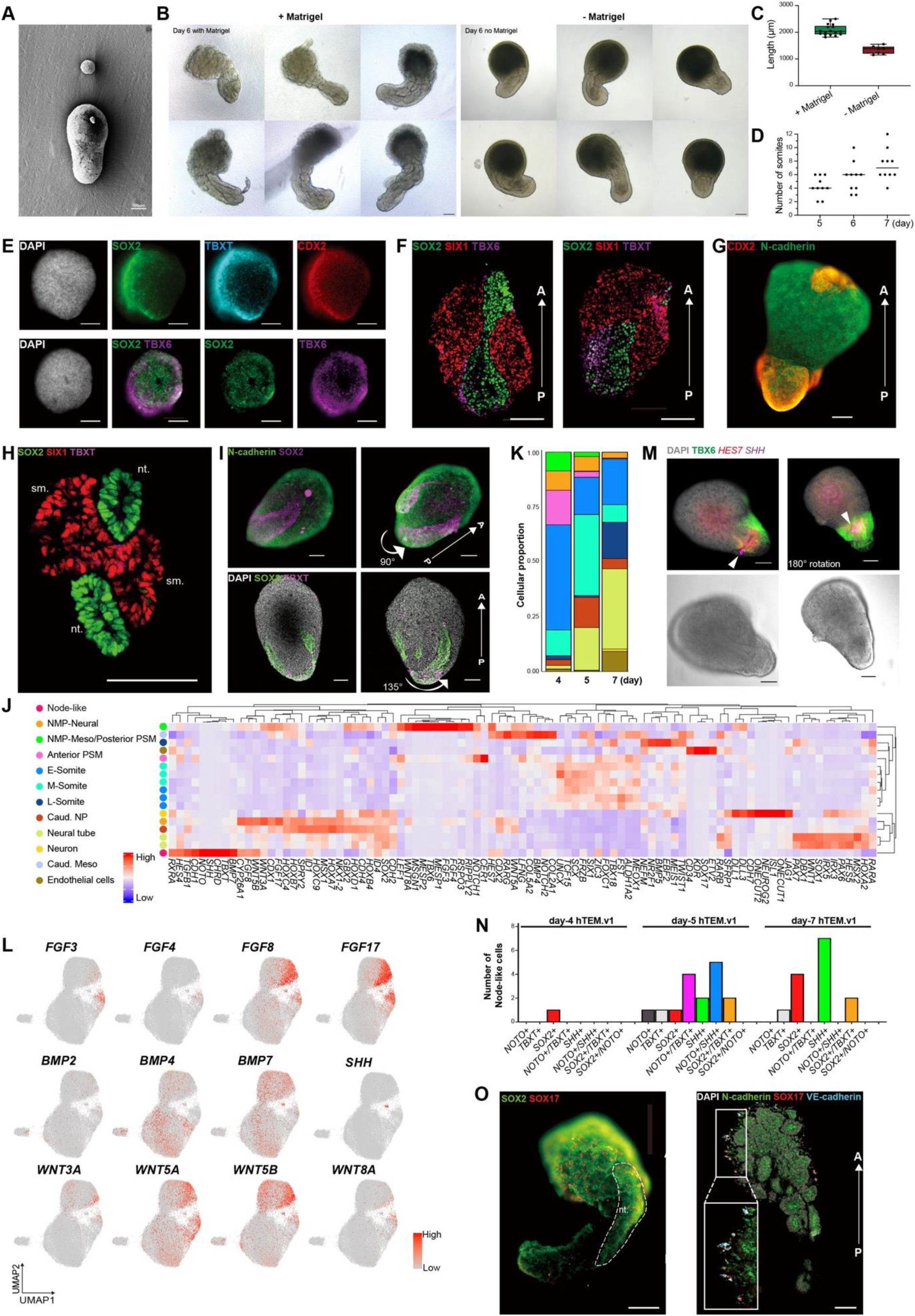
Characterization of hTEM.v1, related to Figure 1. (A) SEM image showing the size contrast between a representative day0-hESC spheroid (top) and day-4 hTEM.v1 (below). (B) Representative images of day-6 hTEM.v1 with or without 4% Matrigel. Scale bars, 200 μm. (C) Box plot showing length measurements of day-6 hTEM.v1 with (n = 15) or without (n = 6) 4% Matrigel. Data are presented as mean ±standard deviation. (D) Number of somite pairs in hTEM.v1 over days 5-7. Each dot represents an individual hTEM.v1 randomly chosen from 3 biological replicates. Vertical lines represent mean values. (E) Top row, 3D projection of day-2 hTEM.v1 showing polarized patterns of bi-NMPs (SOX2, TBXT, CDX2). Bottom row, complementary TBX6+ PSM and SOX2-regions. (F) Immunofluorescence images of longitudinally sectioned day-4 hTEM.v1 showing A-P patterned bi-NMPs (TBXT, SOX2), neural plate (SOX2), PSM (TBX6), and flanking somitic (SIX1) cells. (G) 3D projection of A-P symmetry breaking of day-4 hTEM.v1 with posterior tail bud (CDX2) and anterior somitic cells (N-cadherin). Scale bar, 100 μm. (H) Immunofluorescence image showing the duplication of the neural tubes with flanking somites in transversely sectioned day-7 hTEM.v1. (I) 3D projections of day-4 hTEM.v1 showing the duplication of neural plate (SOX2) structures along A-P axis. TBXT was absent from the presumed ventral side and restricted to the posterior-most region. nt., neural tube. sm., somite. (J) Heatmap of scRNA-seq showing clusters of transcripts associated with cell identities of hTEM.v1 (days 4/5/7) as seen in Figure 1I. (K) Changes in cell type composition in hTEM.v1 over days 4-7. Colors for cell types are in consistent with (J). (L) UMAP showing transcript levels for FGF, BMP and WNT family members in hTEM.v1 (days 4/5/7). (M) HCR-IF images of day-4 hTEM.v1 showing the node-like (*SHH*, arrowhead) structure adjacent to PSM (*HES7*, TBX6). (N) Counts of the node-like cells from hTEM.v1 scRNA-seq data, based on detected marker genes at listed days. (O) Left, 3D projection showing anteriorly positioned vascular endothelial cells (SOX17). Right, Immunofluorescence of vascular endothelial cells (SOX17, VE-cahderin) at the outer layer of epithelialized somites (N-cadherin). All scale bars in the immunofluorescence images are 100 μm.

**Figure S2.**
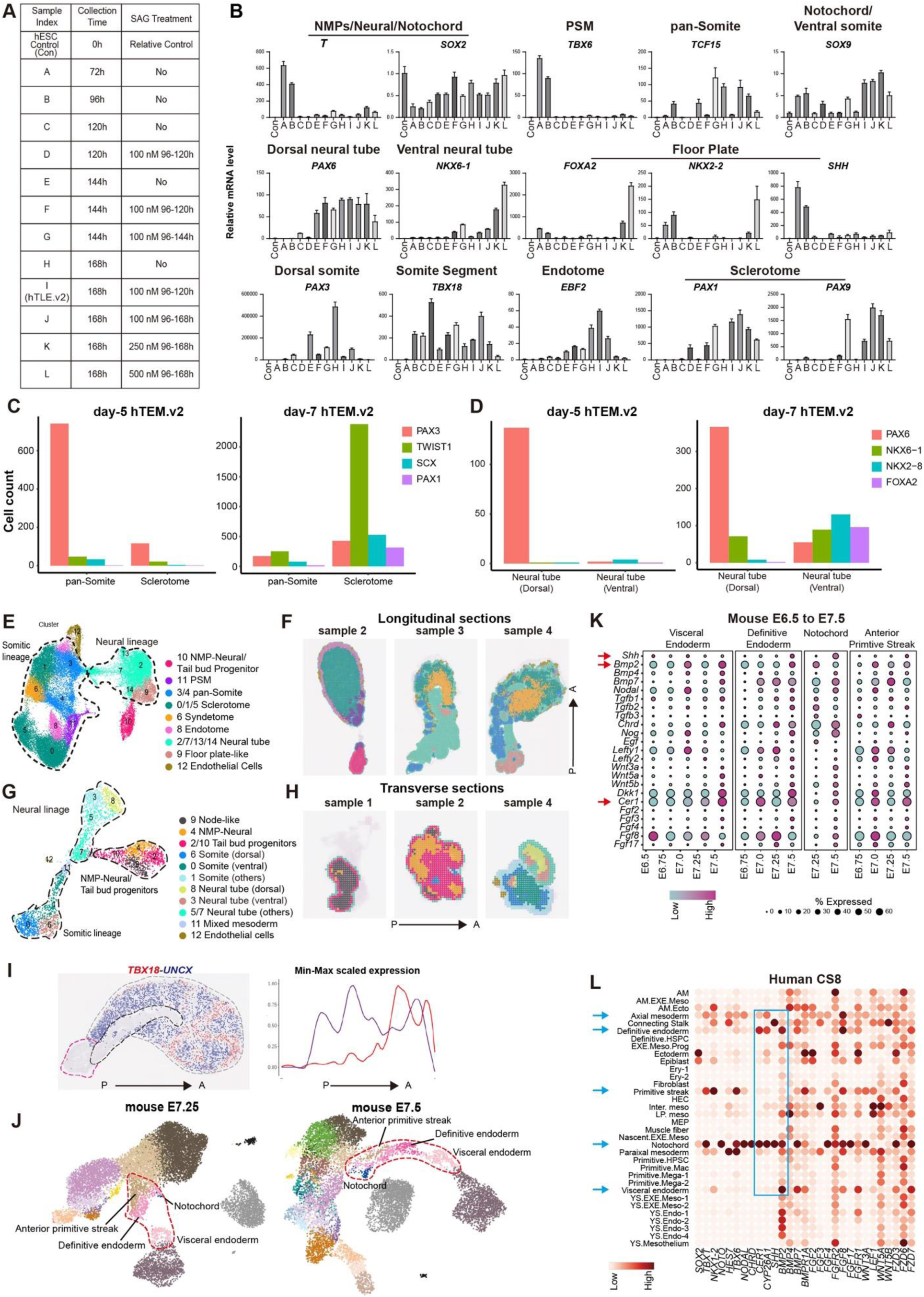
Characterization of hTEM.v2, related to Figure 2. (A) Experimental design for timely and varied SAG concentrations, based on the hTEM.v1 protocol. (B) qPCR showing expression levels of marker genes for indicated cell types. Sample indices are the same as those in (A). Data are presented as mean ±standard deviation. Data were reproduced twice. (C-D) Numbers of D-V patterned cells from somitic (C) and neural (D) clusters in hTEM.v2 scRNA-seq data. Dorsal somite marker; PAX3. Ventral somite markers; TWIST1, SCX, PAX1. Dorsal neural tube marker; PAX6. Ventral neural tube markers; NKX6-1, NKX2-8, FOXA2. (E-F) UMAP (E) and spatial UMAP (F) showing identified cell types from transversely sectioned day-7 hTEM.v2 (n = 4). Transverse samples 1-4 are sections from posterior to anterior positions. (G-H) UMAP (G) and spatial UMAP (H) showing identified cell types from longitudinally sectioned day-7 hTEM.v2 (n = 4). (I) Spatially expressed *TBX18* (anterior) and *UNCX* (posterior) indicating somite segmentation in longitudinal section of day-7hTEM.v2 (Visium HD sample 1). Signals of *TBX18* and *UNCX* were quantified and scaled along the A-P axis. (J) UMAP trajectory showing notochord emergence in E7.25-7.5 mouse embryos (E-MTAB-6967). (K) Dot plot showing the expression of signals emanating from indicated tissues during early mouse embryogenesis (E-MTAB-6967). (L) Dot plot showing the expression of signals emanating from the indicated tissues in the human CS8 embryo (HRA005567). Plot was generated using the online tool in http://cs8.3dembryo.com/.

**Figure S3.**
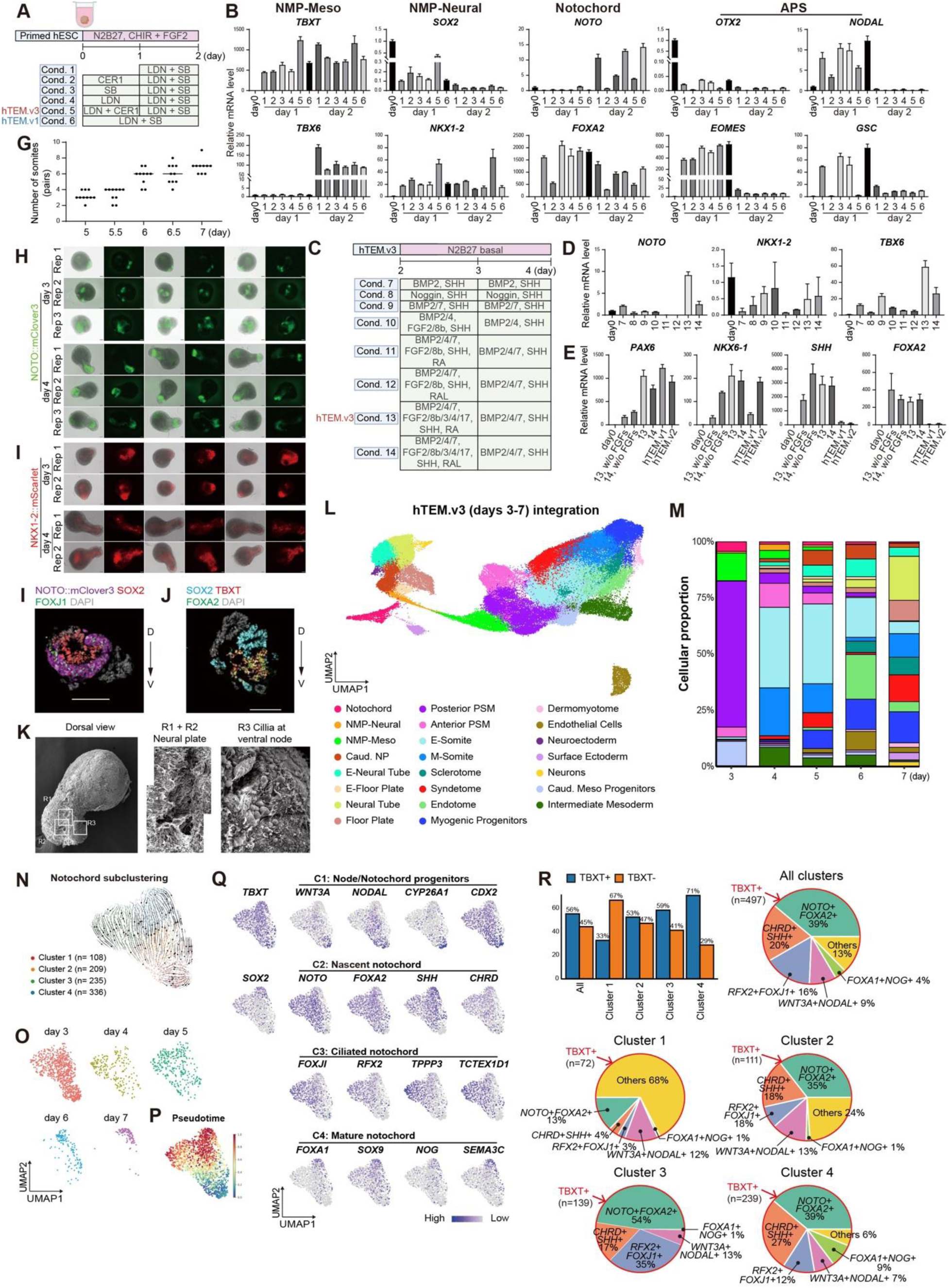
Experimental design, morphological and cellular composition of hTEM.v3, related to Figure 3. (A) Experimental design to test the conditions of timely BMP and NODAL inhibition for notochord cells and bi-NMP induction. (B) qPCR results showing marker levels of bi-NMPs, notochord and anterior primitive streak under test conditions shown in (A). Data are presented as mean ±standard deviation. Results were reproduced more than three times. (C) Experimental design to test combinations of BMPs, FGFs and SHH for concordant maintenance and differentiation of notochord, NMP-Neural and NMP-Meso. (D) qPCR showing expression of indicated lineage markers for notochord (*NOTO*), NMP-Neural (*NKX1-2*), NMP-Meso (*TBX6*) at day 4, from conditions listed in (C). Data are presented as mean ±standard deviation. Results were reproduced more than three times. (E) qPCR results comparing expression levels of dorsal (*PAX6*) and ventral (*NKX6-1*, *SHH*, *FOXA2*) markers in neural and notochord cells in day-6 hTEM.v3. Test conditions are shown in (C). Data are presented as mean ±standard deviation. Results were reproduced more than three times. (F) Dot plot showing numbers of segmented somite pairs in hTEM.v3 at days 5-7. (G) Representative images showing the caudal enrichment of NOTO:mClover3+ notochord cells in the midline of hTEM.v3 at days 3-4. (H) Representative images showing the gradual enrichment of NKX1-2:mScarlet+ caudal neural plate progenitor cells in the midline of hTEM.v3 at days 3-4. (I-J) Immunofluorescence images of transversely sectioned day-5.5 hTEM.v3 exhibiting the bi-layered organization of dorsal neural plate (SOX2) and ventral notochord (TBXT, NOTO, FOXJ1, FOXA2). (K) SEM images of day-4 hTEM.v3 (n = 2); dorsal view. Regions of interest (R1/2/3) were dash lined. Zoomed views of R1+R2 and R3 are displayed, highlighting the neural plate and the ciliated ventral node. (L) UMAP showing identified cell types from hTEM.v3 scRNA-seq integration (days 3-7, total of 100,370 cells). (M) Changes in cell type composition of hTEM.v3 from day 3-7. Colors of cell types correspond to those in (L). (N-P) RNA velocity (N), sub-clustering (O), and pseudotime (P) analysis on notochord cells (n = 888) subset from hTEM.v3 scRNA-seq dataset in (L). (Q) UMAP showing expression of markers for indicated notochord subtypes. (R) Relative proportions of notochord subtypes identified from (N).

**Figure S4.**
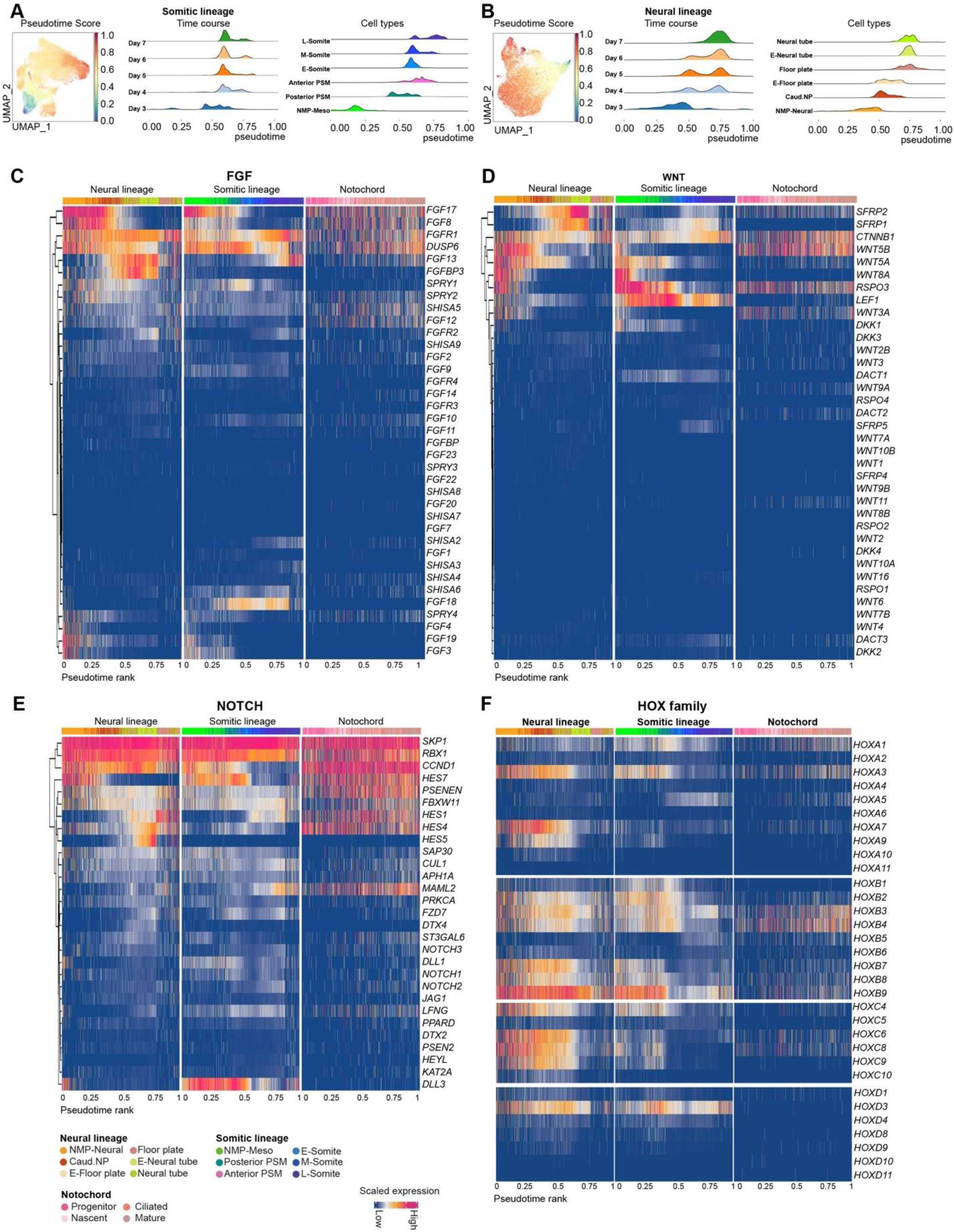
A-P organization of cell types and gene expression profiles inferred from scRNA-seq integration of hTEM.v3 (days 3-7), related to Figure 3. (A-B) Pseudotime analysis showing the progression of somitic (A) and neural lineages (B). Cells analyzed here were subset from the hTEM.v3 (days 3-7) scRNA-seq dataset in Figure S3L. (C-F) Heatmap showing scaled expression of FGF (C), WNT (D), NOTCH (E) pathways and the *HOX* (F) genes along pseudotime inferred A-P axis.

**Figure S5.**
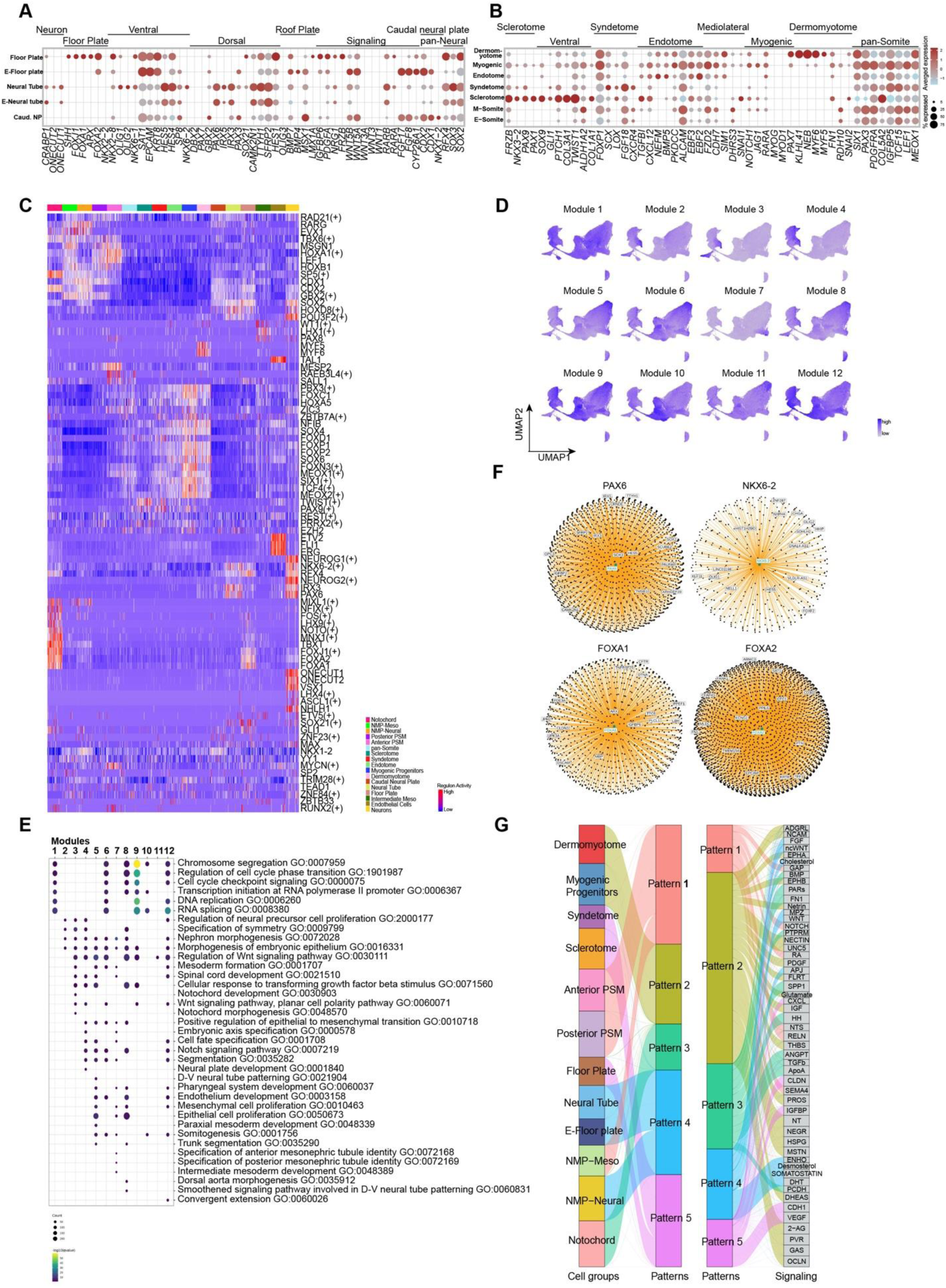
Molecular profiles of hTEM.v3, related to Figure 4. (A-B) Normalized gene expression profiles of listed neural (A) and somitic (B) lineages from hTEM.v3, reflecting the D-V axis formation in hTEM.v3. (C) Heatmap showing regulon activities represented by selected transcription factors (TFs) in 500 randomly sampled cells in hTEM.v3. (D) UMAP showing the regulon module activity over trajectories for hTEM.v3. (E) Gene Ontology (GO) enrichment analysis based on top ranking genes enriched in regulon modules from Figure 4I. (F) Gene regulatory networks of *PAX6*, *NKX6-2*, *FOXA1* and *FOXA2* in hTEM.v3. Top 15 interacted genes were displayed. (G) Cell-cell communication patterns grouped by cell types from hTEM.v3 (days 3-7; left). Significantly enriched signaling pathways belonging to each pattern (right).

**Figure S6.**
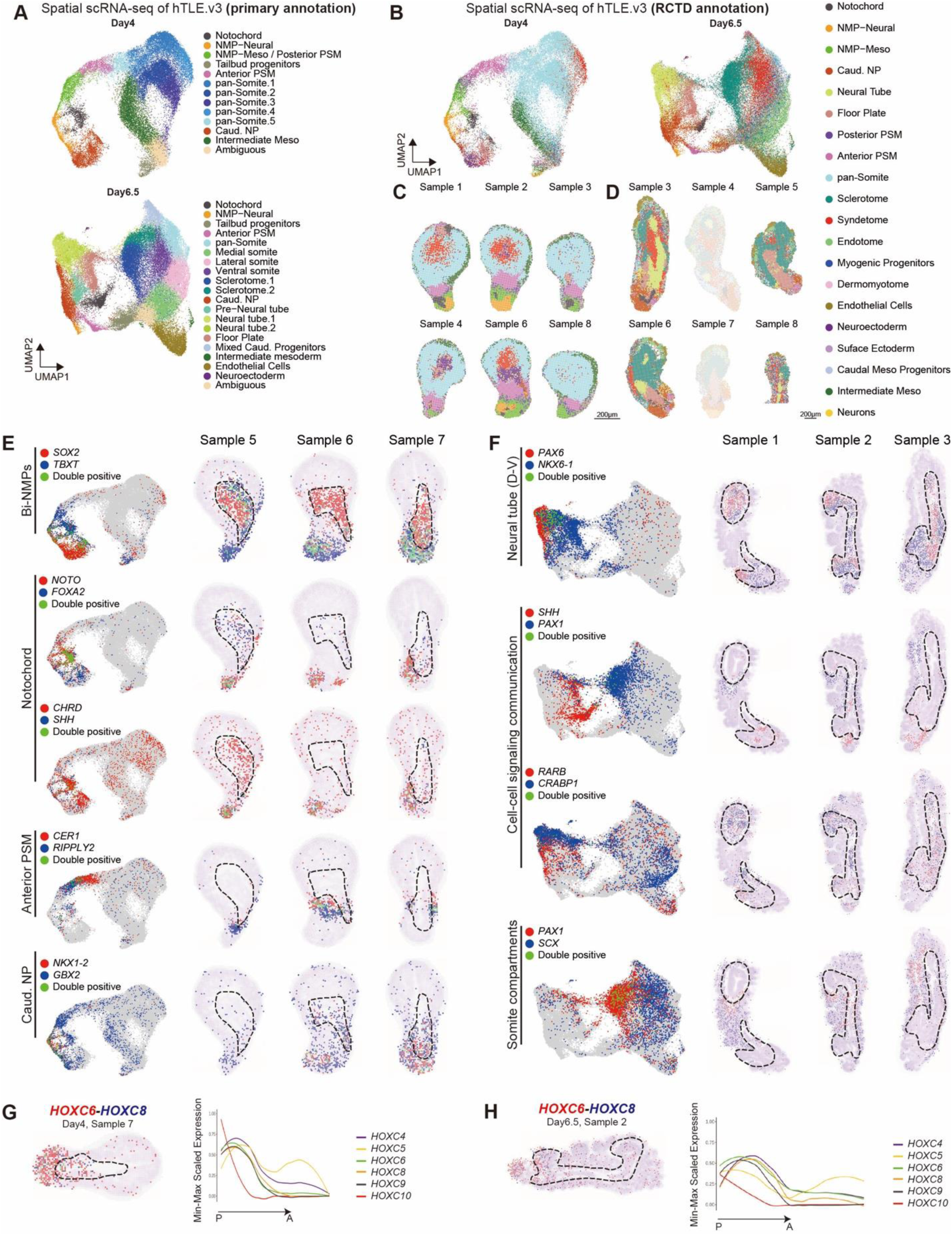
Spatially resolved hTEM.v3 cell type organization using Visium HD, related to Figure 4. (A) UMAP showing the primary cell types identified from Visium HD results of day-4 and day-6.5 hTEM.v3. (B) UMAP showing the RCTD cell types using reference annotations from hTEM.v3 scRNA-seq data in Figure S3L. To reduce annotation complexity and to discern annotation colors in RCTD, ‘E-Somite’ and ‘M-Somite’ from scRNA-seq reference were combined as ‘pan-Somite’ in RCTD. ‘E-Neural tube’ and ‘Neural tube’ from scRNA-seq reference were merged as ‘Neural tube’ in RCTD, ‘E-Floor plate’ and ‘Floor plate’ from scRNA-seq reference were merged as ‘Floor plate’ in RCTD. Colors and annotations are listed at right side. (C-D) Spatial UMAP showing the RCTD annotated cell types from Visium HD resolved from day-4 (C) and day-6.5 (D) hTEM.v3. Colors and annotations are from (B). (E-F) UMAP and spatial expression patterns of indicated genes in longitudinally sectioned day-4 (E) and day-6.5 (F) hTEM.v3. Dashed lines, the neural plate (day 4) and neural tube (day 6.5) structures. (G-H) Spatial expression of *HOXC6* and *HOXC8* at day 4 (G) and day 6.5 (H) in hTEM.v3. Signals of *HOXC* family genes were quantified and scaled along the direction of P-A at the right side.

**Figure S7.**
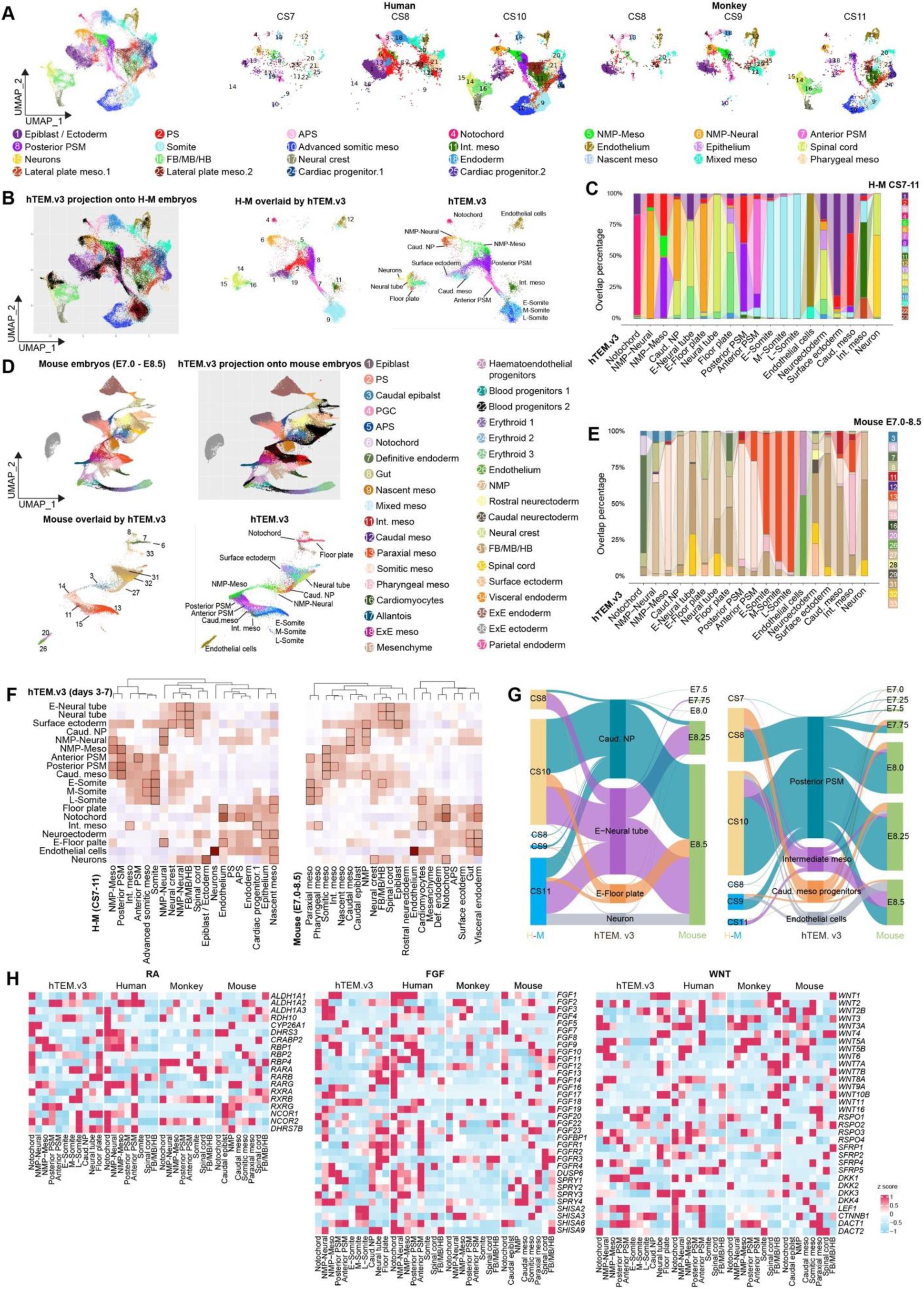
Cross-species comparison between hTEM.v3 and embryos from human, monkey, and mice, related to Figure 5. (A) Ortholog UMAP trajectory of organogenesis in early human (CS7/8/10) and monkey (CS8/9/11) embryos. See ‘Methods’ for published human and monkey embryo scRNA-seq datasets. (B) Left, hTEM.v3 projection onto public human-monkey (H-M) embryo integration dataset. Middle, H-M cell types overlaid by hTEM.v3. Colors and annotations for H-M are consistent with those in (A). Right, hTEM.v3 cell types overlapping with H-M. Colors and annotations for hTEM.v3 are consistent with Figure S3L. (C) Sankey plot showing a detailed cell type overlay between hTEM.v3 and H-M embryos dataset. Colors and annotations for H-M are consistent with those in (A). (D) Top-left, UMAP showing cell types in mouse embryos (E7.0-E8.5). Top-right, UMAP showing hTEM.v3 projection on mouse embryo data (E7.0-E8.5). Bottom-left, mouse cell types overlaid by hTEM.v3. Bottom-right, hTEM.v3 cell types overlapping with mouse embryos. Colors and annotations for hTEM.v3 are consistent with Figure S3L. (E) Sankey plot showing the detailed cell type overlay between hTEM.v3 and mouse embryo dataset. Colors and annotations for mouse embryos are consistent with those in (D). (F) Heatmap showing cluster similarities in listed cell types between hTEM.v3, H-M, and mouse embryos, respectively. The highest correlations are highlighted by black boxes. The second highest is in red box. (G) Sankey diagram showing relationships between selected cell types in hTEM.v3 with human (CS7/8/10), monkey (CS8/9/11) and mouse embryos (E7.0-E8.5). (H) Heatmap showing the scaled expression pattern of genes involved in RA, FGF and WNT signaling pathways in indicated cell types across datasets of hTEM.v3 and embryos.

**Figure S8.**
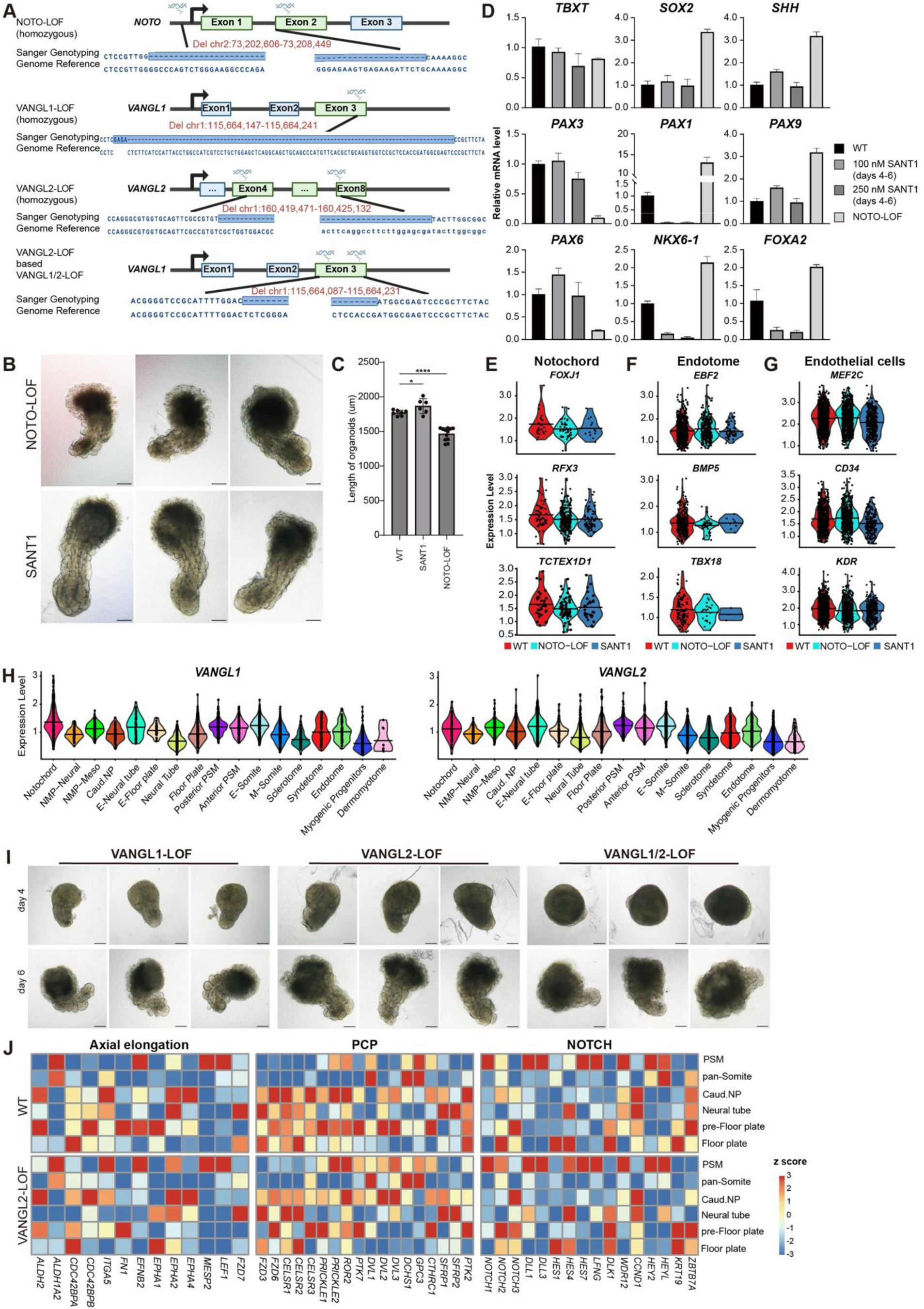
Morphological and molecular changes upon genetic and chemical perturbations in hTEM.v3, related to Figure 6. (A) Schematic showing the CRISPR/Cas9 design and genotyping validations of NOTO-LOF, VANGL1-LOF, VANGL2-LOF, and VANGL1/2-LOF hESC lines. (B) Representative images of day-6 hTEM.v3 in NOTO-LOF or, SANT1 treatment (250 nM, days 4-6). Scale bars, 200 μm. (C) Comparison of length measurements of day-6 hTEM.v3 between WT, NOTO-LOF and SANT1 treatment (250 nM, days 4-6) (n = 6-8 each group). \**p* value < 0.05 and \*\*\*\**p* value < 0.0001 was calculated by Student’s t test. n.s., no significance. The results were reproduced in more than 3 biological replicates. (D) qPCR showing changes in D-V patterning in WT, NOTO-LOF and SANT1 treatments (100 nM or 250 nM at days 4-6) in day-6 hTEM.v3. Data are presented as mean ±standard deviation. Results were reproduced twice. (E-G) Violin plot showing changes in expression of genes related to notochord, endotome and endothelial cells in NOTO-LOF or, SANT1 treatment (250 nM, days 4-6). (H) Violin plot showing expression of *VANGL1* and *VANGL2* across listed cell types in hTEM.v3 dataset from Figure S3L. (I) Representative bright field images of day-6 hTEM.v3 in VANGL1-LOF, VANGL2-LOF, or VANGL1/2-LOF. Scale bars, 200 μm. (J) Scaled expression levels of key genes involved in axial elongation, PCP and NOTCH signaling in day-6 hTEM.v3 dataset from Figure 6A.

## Methods

### Culture of human ES and iPS cells

The following human ESC and iPSC lines were used: wild type H9-hESC (Sex: female, WiCell, WAe009-A), K3-iPSC (Sex: male)^108^ generated from human neonatal foreskin fibroblasts (ATCC, PCS-201-010), H9-hESC line carrying IRES-mClover3 allele in the 3’ UTR of *PAX3* and IRES-mScarlet allele in the 3’ UTR of *EBF2 (*PAX3:mClover3; EBF2:mScarlet dual reporter), H9-hESC line carrying IRES-mScarlet allele in the 3’ UTR of *NKX1-2* (NKX1-2:mScarlet reporter), H9-hESC line carrying IRES-mClover3 allele in the 3’ UTR of *PAX3* and IRES-mScarlet allele in the 3’ UTR of *NOTO (*NOTO:mClover3 reporter), H9-hESC line carrying IRES-mClover3 allele in the 3’ UTR of *MYF5 (*MYF5:mClover3 reporter), and H9-hESC derived NOTO-LOF, VANGL1-LOF, VANGL2-LOF, and VANGL1/2-LOF (Figure S8A). H9-hESC lines and K3-iPSCs were routinely cultured and passaged as described previously.^109,110^ Briefly, 5 × 10^4^/cm^2^ cells were seeded onto culture-treated petri dishes coated with 1:200 diluted Geltrex (Thermo, A1413302). The basic culture media is in-house prepared using DMEM/F-12 w/o glutamine (Thermo, 21331020 or Servicebio, G4514), supplemented with 0.5% Probumin (Sigma, 821001), 1x Antibiotic-Antimycotic (Thermo, 15240062), 1x MEM NEAA (Thermo, 11140050), 1x Trace Elements A/B/C (Corning), 64 μg/mL Ascorbic acid magnesium (TCI, A2521), 10 μg/mL Transferrin (Athens Research and Technology), and 1 x GlutaMax (Thermo, 35050061). To maintain pluripotency of hESCs and hiPSCs, the basic culture media is completed by addition of 10 ng/mL Heregulin β1 (Qkine, QK045), 10 ng/mL Activin A (Qkine, QK001), 8 ng/mL FGF2-G3 (145aa) (Qkine, QK052) and 200 ng/mL IGF-1 LR3 (Qkine, QK041). The complete pluripotency maintenance medium is abbreviated as HAIF. hESC lines and K3-iPSCs were cultured in HAIF with media changes every 24 hours at 37 °C in 5% CO2 and passaged at 90% confluency using Accutase (Thermo, 00-4555-56).

### Generation of CRISPR/Cas9 knock-in fluorescent reporter H9-hESC lines

To generate knock-in fluorescent reporter lines from H9-hESCs, we utilized the homology directed repair approach based on Cas9/gRNA introduced double strand break. The px330 vector expressing human codon-optimized Cas9 and sgRNA was obtained from Addgene. Oligos for sgRNA targets were individually cloned into px330 using restriction enzyme BbsI. Homology directed repair (HDR) donor arms flanking the internal sequences of ribosome entry site (IRES) and fluorescent protein coding sequence (mClover3 or mScarlet) was synthesized and cloned into a pUC57 vector by BGI. The px330 with gRNAs targeting the gene of interest, homology donor arm vector and corresponding surrogate reporter vector (pRGS, PNA Bio) (vector ratio of 2:3:1, total of 9 μg vectors per 3 x 10^6^ cells) were electroporated into H9-hESCs using a Neon electroporation system (voltage = 1050 V, width = 30 ms, pulse = 2 cycles) (Thermo, MPK10025). To enhance cell viability, 1x CEPT^111^ (in-house preparation) was used during electroporation and FACS sorting. Two days after electroporation, single cells were isolated by FACS sorted and plated on pMEF (Sigma, PMEF-NL-P1) coated 96-well plates based on pRGS surrogate reporter signals. Single cell derived clones were screened by genotyping PCR, expanded in HAIF and subject to hTEM protocols. To generate CRISPR/Cas9 knock-out H9-hESC lines, gRNAs targeting upstream and downstream regions of exon(s) were used together with a surrogate reporter vector. Electroporation and single cell clone screening procedures were the same as generating knock-in cell lines. All oligos for sgRNA targets are listed in the Table S1.

### Generation of human trunk embryoid models (hTEMs)

Routinely 2D passaged hPSCs (H9-hESC lines and K3-iPSC) at 80% confluency were dissociated using Accutase for 5 min at 37 °C and counted by a hemocytometer. To start 3D spheroid formation, a total of 1-2 million cells in 5 mL of HAIF medium with 10 μM Y-27632 (Aladdin) were seeded per well of an ultra-low attachment 6-well plate (Thermo, 174929). The plate was then placed on an orbital shaker (Thermo, 88881104) at 110-120 rpm at 37 °C in 5% CO2. The next day, a media change consisting of HAIF with 10 μM Y-27632 was applied. hPSC spheroids were allowed to form at size of 200-220 μm in diameter within 40 hours of shaking. To generate hTEMs, hPSC spheroids were collected with a wide-bore 1 ml tip and transferred into a 1.5 mL Eppendorf tube, then washed with N2B27 basal medium. N2B27 basal medium comprises a 1:1 mix of DMEM/F12 and Neurobasal A (Thermo, 21103049) supplemented with 1×B27 (Thermo, 17504001), 1× N2 (Thermo, 17502048), 1.5× GlutaMAX, 1×MEM NEAA, 1x Sodium Pyruvate (Thermo, 11360039), 1x Antibiotic-Antimycotic, and 64 μg/mL Ascorbic Acid Magnesium. The HAIF primed hPSC spheroids (day 0 of hTEM) were then individually transferred to a well of an ultra-low attachment 96 well plates (Thermo, 174929), subject to hTEM protocols.

For hTEM.v1, hPSC spheroids (day 0) were induced by culture in bi-NMP induction media composed of N2B27 basal media and supplements of 10 μM CHIR-99021 (CHIR) (MCE, HY10182), 500 nM LDN-193189 (LDN) (Tocris, 1517128), 10 μM SB-431542 (SB) (Aladdin, S125924), and 20 ng/mL FGF2-G3. 48 hours later, N2B27 basal media was replenished. At day 4, media is replaced by N2B27 basal media supplemented with 4% Geltrex (Thermo, A1413302) of Matrigel (Mogengel, 827775) and 1 μM all-trans retinal (RAL) (Aladdin, A122355) to support somitogenesis and neural tube elongation.

For hTEM.v2, 100 nM SAG (Aladdin, S872455) was added at day 4-5 based on the hTEM.v1 protocol, followed by replenishment of N2B27 basal medium containing 4% Matrigel and 1 μM RAL at day 5.

For hTEM.v3, the hTEM.v1 day 0-2 protocol is modified by replacing SB with 20 ng/ml human recombinant CER1 (MCE, HY-P7822) for the first 24 hours. At day 2-3, media was replenished with N2B27 basal medium supplemented with 0.2 ng/mL SHH (R&D, 8908-SH), 2 ng/mL BMP4 (R&D, 314-BP), 1 ng/ml BMP2 (Qkine, QK007), 0.5 ng/ml BMP7 (R&D, 354-BP), 8ng/ml FGF2-G3, 4 ng/ml FGF3/4/8b/17 (MCE, HY-P700065/HY-P7014/HY-P70533/HY-P700060), 1 nM all-trans retinoic acid (RA) (Aladdin, R106320) and 0.2% Matrigel. At day 3-4, medium was changed to N2B27 basal medium supplemented with 0.4 ng/ml SHH, 0.5 ng/ml BMP2/4/7, 10 ng/ml WNT5A (R&D, 645-WN) and 0.4% Matrigel. At days 4-7, medium was replaced with N2B27 medium containing 4% Matrigel and 200 nM RAL. 10 ng/ml WNT3A (R&D, 5036-WN) and 0.2 ng/ml Heregulin β1 were included at days 4-7 to enhance neural tube genesis.

All hTEM cultures were limited to 7 days due to accumulated cell death in the anterior region and ceased axial elongation.

### Inclusion criteria of hTEM embryoids

All hTEM embryoids were collected using wide-bore tips from ultra-low attachment 96 well plates within 7 days of culture. Embryoids were inspected under a phase contrast microscope, based on morphometric features resembling human CS8-10 embryos. At day 3-4 (equivalent to CS8), embryoids undergo uniaxial symmetry breaking and become oval shaped. Embryoids with cylindrical morphology and visible mediolateral narrowing near the caudal end were selected for further use. The neural plate is to be visible as a groove along the midline extending from the posterior end and flanked by a shaded area anterior to the tailbud, reflecting the PSM cells (Figures 1B and 3B). If using the NOTO:mClover3 or NKX1-2:mScarlet reporter line, polarized and caudal accumulation of NOTO+ or NKX1-2+ cells at day 3 and axial distribution of these cells in the midline at days 4-5 (Figures S3F and S3G) were expected. This is equivalent to the C&E movements of caudal trunk progenitors in natural CS8 human embryos. Embryoids with a correct body plan require culture in media supplemented with 4% Matrigel over days 4-7 to allow somitogenesis and neural tube morphogenesis. At days 4-7 (equivalent to CS9 to CS10) the following criteria were applied: (1) somites were to be clearly segmented after day 5; (2) presence of a lumen representing the closed neural tube was observed along the midline from hTEMs (Figures 1B, 2B, and 3B); (3) a single body axis with less-dense tailbud cell populations on the posterior end and high-dense unsegmented somite cells surrounding the neural tube tip on the anterior end. At day 4, ∼70% of the embryoids with a correct body plan were collected for analysis or subject to further Matrigel culture. At days 5-7, ∼30% of the hTEM.v1/2 and ∼20% of hTEM.v3 satisfied the criteria specified above.

### Morphometric feature measurements

The length of hTEMs were analyzed using ImageJ. To calibrate the length in pixels, a standard ruler is utilized to set the scale of images with the same magnification. The anterior-most and posterior-most points of the embryoids were set as start and end points for measurements. The length was measured along a custom defined midline along the direction of axial elongation. The counting of somite pairs was based on their order from the posterior end. The presence of two rows of somite segments flanking a neural tube structure was confirmed before counting all somite pairs along the A-P axis.

### RNA extraction & RT-qPCR analysis

hTEMs RNA was extracted using E.Z.N.A. MicroElute Total RNA Kit (OMEGA, R6831-02). 500 ng of total RNA was reverse transcribed into cDNA using iScript Reverse Transcription Supermix (Bio-rad, 1708841) according to manufacturer’s instructions.

Quantitative real-time PCR was performed using Taq Pro HS Universal Probe Master Mix (Vazyme, QN113-01) on QuantStudio 7 Pro Real-Time PCR System (Applied Biosystems, A43183). Expression levels for each gene re normalized to *18S rRNA* check italics as an endogenous control using the ^ΔΔ^Ct method. Taqman probes used in this study are listed in the Table S1.

### Scanning electron microscopy (SEM)

Embryoids were washed with DPBS to remove Matrigel, then fixed with 4% PFA for 30 minutes at room temperature. Next, samples were treated with 1% osmium tetroxide in distilled water for 1 hour and then washed three times with distilled water for 10 minutes each. Samples were then dehydrated by serial washes in 70% to 100% ethanol. Critical point drying (CPD) was carried out using a Tourimis Samdri (PVT30) Critical Point Dryer. To prepare samples for imaging, an 80:20 platinum/palladium sputtering process was conducted with rotation, depositing a 3 nm conductive layer using the Quorum Q150T Automatic Coating System. Finally, SEM images were acquired using a Hitachi SU8010 cold-field emission scanning electron microscope at 9.8 kV.

### Whole-mount hTEM immunostaining and imaging

hTEM embryoids were transferred to 1.5 ml Eppendorf tubes, fixed in 4% paraformaldehyde (PFA)/DPBS at RT for 1 hr and then washed with 1x DPBS at 10 min intervals for 30 min to remove residual PFA. For antigen retrieval, samples were immersed in warm 0.5% SDS/DPBS (preheated at 55 °C) for 15 mins, followed by incubation with 0.5% Trition X-100 in DPBS for 30 min. After blocking samples with Duolink blocking solution (Sigma) for 30 min at room temperature, samples were incubated with primary antibodies, diluted in MAXbind staining medium (Active Motif) overnight at 4 °C. Next, samples were washed three times with 0.04% Tween-20/DPBS (5 min each) and then incubated with DAPI (TCI, 1ug/ml) and Alexa Fluor secondary antibodies (Thermo, 1:500 dilution) in MAXbind staining medium for 2 h at room temperature. This was followed by one wash in 1 ml MAXwash washing medium (Active Motif) and two washes in 0.04 % Tween-20/DPBS (5 mins each) at room temperature. Optionally, a clearing procedure with Optimus Clearing Solution was used.^112^ Finally, samples were individually transferred into chamber slides (iBidi, 81811), cured by ProLong Gold antifade mountant (Thermo, P36934) overnight before imaging. All samples were imaged and analysed using a BZ-X All-in-one inverted fluorescent microscope (Keyence) or SP8 inverted confocal microscope (Leica). Antibodies used in this study are listed in the Key Resources Table.

### Cryosectioning of hTEMs

hTEM embryoids were washed twice with ice-cold DPBS to remove Matrigel and fixed by 4% PFA, then dehydrated in 30% sucrose (Aladdin, S112234) overnight at 4°C. Samples were then transferred into a 1cm x 1cm cryomold, positioned and embedded in OCT compound (Tissue-tek, 4583). OCT embedded molds were snap frozen in dry ice and 95% isopentane (Macklin, I813377) and transferred to -80°C overnight before sectioning. The frozen and fixed samples were sectioned using a cryostat microtome (Leica, CM1950) at 10 μm. Sections were kept at -20°C before further analysis. All buffers in contact with hTEM embryoids were pre-treated with RNaseOUT (Thermo, 10777019) to preserve RNA integrity for Visium HD process.

### Time-lapse imaging of hTEMs

Bright-field and fluorescent images of live hTEMs were taken with a BZ-X All-in-one inverted fluorescent microscope (Keyence) in the ‘Time lapse capture’ mode using a 10× plan objective. Images were captured using the z-stack mode at a step depth of 6-10 μm, spanning a total of ∼100 μm. For time-lapse imaging, the incubator module was set at 37 °C and 5% CO2 and images taken every 30 min or 60 min. To generate the time-lapsed video, stacked snapshots within the best focus range from each time point were used.

### In situ hybridization chain reaction (HCR)

hTEM embryoids were fixed in 4% PFA for 30 min at room temperature (RT), followed by one wash with ice-cold DPBS. Before HCR, fixed samples were pre-treated with 0.5% SDS/DPBS for 15 min at RT, washed twice with 0.5% Triton X-100/DPBS and washed three times in 0.1% Tween-20/DPBS for 5 min each wash. HCR was performed following manual instructions (Molecular Instruments). In brief, samples were incubated in hybridisation buffer (HB) for 5 min at RT, then for 30 min at 37 °C. Probes were prepared at 8 nM in HB and incubated for 30 min at 37 °C before use. Samples were then incubated with probes for 12-16h on a thermal cycler (Bio-rad) at 37 °C. The next day, samples were washed with probe wash buffer (WB) three times for 15 min each at 37 °C, then with 5x SSCT (5x SSC and 0.1% Tween-20 diluted in UltraPure Water) three times for 15 min each at RT. Samples were then pre-amplified in probe amplification buffer for >30 min at RT. Amplifier hairpins (h1 and h2) were prepared by heating at 95 °C for 90 sec separately followed by cooling down for 30 min at RT in dark. h1 and h2 were then mixed at 6 nM in amplification buffer and incubated with samples for 12-16 hrs at RT in the dark. Before imaging, samples were washed three times in SSCT for 15 min, and stained with DAPI. All HCR images were acquired and processed with a Nikon Ti2E inverted microscope. All HCR probes with associated hairpins are listed in the Table S1.

### Single-cell RNA-seq

hTEMs (hTEM.v1; day3 = 35, day 4, n = 31, day 5, n = 22, day7, n = 11. hTEM.v2; day 5, n = 22, day 7, n = 16; hTEM.v3; day 3, n = 35, day 4, n = 23, day 5, n = 12, day 6, n = 9, day 7, n =14. NOTO-LOF; hTEM.v3 day 6, n = 11. SANT-1 (250 nM at days 4-6) hTEM.v3 day 6, n = 8. VANGL2-LOF; hTEM.v3 day 6, n =18.) were washed with N2B27 basal medium twice and then dissociated using 1:1 TrypLE (Thermo) and Accutase at 37 °C for 10-15 min. Single cells were counted and adjusted to 10^6^/ml in 1 ml N2B27 basal medium, filtered through a 40 μm cell strainer and loaded onto Chromium Single Cell 3’ Library and Gel Bead Kit v3.1 or v4 (10× Genomics). Following the cDNA-amplification reaction, quality control and quantification was performed on the Agilent 4200 Tapestation using the High Sensitivity D5000 kit (Agilent Technologies). Illumina sequencing libraries were constructed by fragmentation, end repair, A-tailing and double-sided size selection, adaptor ligation and sample-index PCR. Quality control and quantification of final libraries were performed on the Agilent 4200 Tapestation using the D1000 kit (Agilent Technologies) and Qubit 4 (Thermo Fisher Scientific). Libraries were then sequenced on NextSeq 2000 (Illumina) with a customized sequencing run format until sufficient saturation was reached.

### Pre-processing of scRNA-seq reads

#### Single-cell RNA-seq data generated in this study

All sequencing reads in this study were mapped to the reference genome GRCh38 using cellranger (v9.0.1) with default parameters. Quality control and downstream analysis were performed with in R (v4.3.3) with Seurat (v4.3.0.1). For each dataset, cells with fewer than 200 genes expressed or > 5% expressed mitochondrial genes were removed. Doublets wre identified and removed using DoubletFinder (v2.0.4) implemented in scutilsR (v0.1.1). Next, ambient RNA contamination was estimated, counts were corrected using celda (v1.18.2) implemented in scutilsR (v0.1.1). Cells with less than 200 genes expressed after count correction were filtered out. Cell cycle phase scores were estimated using the corrected counts and these features were regressed out prior to dimension reduction and UMAP construction using Seurat.

#### Public Single-cell RNA-seq data

Public and pre-processed datasets from E-MTAB-6967 (human CS7), HRA005567 (human CS8), GSE155121 (human CS10), GSE193007 (Cynomolgus monkey CS8/9/11), E-MTAB-6967 (E7.0 to E8.5 with all known lineages within this period) were downloaded and included for cross-species analysis. Data from human and monkey embryos were integrated based on ortholog using biomaRt (v2.62.0) and Seurat’s default integration pipeline. Cell-cycle genes were regressed out using the matrix generated by ScaleData function, followed by dimension reduction, UMAP visualization, and clustering. Cells belonging to extra-embryonic tissues were excluded from downstream analysis to reduce the complexity of annotation and data visualization

#### Dataset integration and batch effect correction

To analyze scRNA-seq datasets created in this study and public H-M embryo data generated on different platforms, Seurat, BBKNN or Harmony were used for integration and batch effect correction, based on best performance. Each dataset from different experiments (this study or public) is considered a batch and contains at least one shared cell type. Briefly, log-normalized and scaled matrices from each dataset were integrated using the best-performed approaches. Harmony (v1.0.1) was used for hTEM.v1 (days 4/5/7) and hTEM.v2 (days 5/7) data integration. BBKNN (v1.1.1) was used for hTEM.v3 (days 3-7) data integration. The reciprocal PCA from Seurat was used to integrate ortholog human (CS7/8/10) and monkey (CS8/9/11) embryo datasets. The integration performance is justified given that the gene-expression profiles are from well-studied cell populations that closely match with the identified cell type datasets reported here.

### RNA velocity analysis

For RNA velocity analysis, the raw fastq sequence data using scvelo (v0.2.5) were firstly reanalyzed to obtain the count matrices containing the spliced and unspliced reads, followed by filtering out cells not subjected to UMAP projection and clustering analysis. Then, RNA velocity was analyzed with velocyto (v0.17.17) in the Python 3.7 environment. Parameters were min_shared_counts= 50 and n_top_genes= 2000 for scv.pp. filter_and normalized function, n_pcs= 30 and n_neighbors= 30 for scv.pp. moments function. Mode was set to be stochastic when computing velocities. The velocity was projected to the UMAP generated previously.

### Pseudotime analysis

Before pseudotime analysis, the scaled matrix in Seurat was converted to the h5ad format using coverFormat function in sceasy (v0.0.7) package. Palantir (v1.3.3) was then used with default parameters in python (v3.9). Note that markers (*NKX1-2* for neural, *TBX6* for somitic and *NODAL* for notochord lineage) for differentiation initiation of each lineage were predetermined for each lineage. The resulting pseudotime scores were displayed based on UMAP embeddings from sub-clustering of each lineage to elucidate developmental dynamics. The A-P axis inference is based on transcriptomic profile of each lineage through time and cell fate specifications in a pseduotime ranked manner.

### SCENIC analysis

SCENIC (v1.3.1) analysis was performed on the integrated hTEM.v3 (days 3-7) scRNA-seq data using R (v4.3.3) and Python (v3.13.2) with arboreto (v0.1.6). Firstly, SCENIC was performed based on motif annotations (hgnc v9) and cisTarget resources of human ‘hg38_refseq-r80_500bp_up_and_100bp_down_tss.mc9nr.feather’ and ‘hg38_refseq-r80_10kb_up_and_down_tss.mc9nr.feather’ following default pipeline in https://htmlpreview.github.io/?https://github.com/aertslab/SCENIC/blob/master/inst/doc/SCENIC_Running.html. Next, the predicated regulon activity scores (RAS) represented by major transcription factors were associated to cell types of hTEM.v3 and grouped by hierarchical clustering to generate the regulon module enrichment heatmap. To this heatmap, correlation matrix was calculated using scaled cell type-associated RAS and clustered by using ‘method = pearson’ from clusterProfiler (v4.14.4) in R. Genes from each regulon module were subject to Gene Ontology analysis and visualized using igraph (v2.0.3) in R.

### Cell-cell communication analysis

CellChat (v2) was used to analyze intercellular communication following cluster annotation. The normalized gene expression matrix from the Seurat object was supplied as input and processed with default parameters. The CellChatDB. human database was used to infer the probability of cell-cell communication with “type = “truncatedMean”, trim = 0.1”. Significant ligand–receptor pairs (p < 0.05) were identified and assigned to signaling pathways. For visualization of overall communication patterns, the summed expression profiles of ligands and receptors were integrated with the predicted communication probability.

### Cross-species cell type projection and comparative analysis

Single-cell transcriptomic datasets were obtained from CS8-CS11 cynomolgus monkey embryos and E7.0-E8.5 mouse embryos, together with the corresponding cell annotations. Gene symbols from monkey and mouse were converted to their human orthologues using the biomaRt package. To map hTEM.v3 cells onto UMAPs of H-M and mouse embryos, cells of hTEM.v3 were down-sampled to 20,000 and then projected using reference mapping approach described in Seurat pipeline https://satijalab.org/seurat/articles/integration_mapping. Briefly, the conserved label anchors between selected objects were identified using FindTransferAnchors function, followed by MapQuery function. The overlapped cell types between objects were determined by using TransferData function. Cross-species cell type projection was subsequently visualized in UMAP and Sankey plot to assess cell type conservation among datasets. ClusterSimilarity was used to determine the pairwise correlation co-efficient across cell types from hTEM.v3 and embryos.

### Preparation of Visium HD spatial gene expression assay

PFA fixed and OCT frozen sectioning (hTEM.v2 and hTEM.v3) slides were prepared as described in the previous section. Sectioning follows the Visium HD Fixed Frozen Tissue Preparation Handbook (10x Genomics, CG000764) for the workflows.

Sequencing was conducted by the Single Cell & Spatial Omics Core, School of Biomedical Sciences at The Chinese University of Hong Kong (https://www3.sbs.cuhk.edu.hk/en/core_laboratories/single-cell-omics-core/.).

### Processing of Visium HD data and visualization

Raw hTEM.v2 and hTEM.v3 Visium HD sequencing reads were mapped to the reference genome using spaceranger (v3.1.3) with default settings then, processed and analyzed with Seurat (5.2.1) running on R (4.4.2). For quality control, spots containing less than 700 total UMI counts were filtered out, as these spots represent areas outside of the actual tissue. Similarly, low-quality spots containing less than 200 total UMI counts and spots containing > 5% mitochondrial counts were filtered out. Following quality control, 3091 spots of transversely sectioned and 39212 spots of longitudinally sectioned day-7 hTEM.v2 were retained with a median transcript read out of 4180 and 2122 unique molecular identifiers (UMI), respectively. For the spatial transcriptome of hTEM.v3, a total of 34,884 (median UMI 2,360) and 58,646 (median UMI 1,260) spots were retained from day 4 and day 6.5 sections, respectively.

Filtered read counts were then processed following the standard Seurat protocol (https://satijalab.org/seurat/articles/visiumhd_analysis_vignette). Briefly, samples were normalized and scaled, followed by dimension reduction using PCA and UMAP. The FindNeighbors function was followed by RunUMAP function with 50 dimensions. The primary clusterings of Visium HD data were identified via the FindClusters function for crude cell type annotations. Unsupervised clustering was conducted using a bin size of 8 μm and Seurat default settings. All other parameters used for each function were kept at default values.

To deconvolute the spot-level data for accurate cell type annotations, the Robust Cell Type Decomposition (RCTD) approach from Seurat was applied using the reference cell types from integrated hTEM.v3 (days 3-7) scRNA-seq data. RCTD clusters were annotated by leveraging the cell type reference in Figure S3L. To visualize spatial UMAP of marker gene expression, customized scripts were devised using SpatialDimPlot function in Seurat for displaying weight normalized read counts listed in Figure 4L-4N, 6G, S5E-S5H.

### Quantification and statistical analysis

All statistical results and graphs were generated by Graphpad Prism. The numbers of samples and types of statistical analyses are given in the figure captions and results sections.

