## Supplementary material for "Human trunk embryoids with patterned anterior-posterior and dorsal-ventral body axes: utility for understanding human development and disease": Key Resources Table

| REAGENT or RESOURCE | SOURCE | IDENTIFIER |
| --- | --- | --- |
| Antibodies | | |
| Mouse anti-SOX2 (2.5 µg/mL) | R&D | Cat# MAB2018; RRID: AB_358009 |
| Rabbit anti-SOX2 (1:200) | CST | Cat# 3579; RRID: AB_2195767 |
| Goat anti-SOX2 (1:100) | Santa Cruz | Cat# sc-17320; RRID: AB_2286684 |
| Rabbit anti-TBXT (1:100) | Santa Cruz | Cat# sc-20109; RRID: AB_2255702 |
| Goat anti-TBXT (2.5 µg/mL) | R&D | Cat# AF2085; RRID: AB_2200235 |
| Goat anti-TBX6 (2.5 µg/mL) | R&D | Cat# AF4744; RRID: AB_2200834 |
| Rabbit anti-SIX1 (1:200) | CST | Cat# 12891S; RRID: AB_2753209 |
| Mouse anti-PAX3 (1:50) | R&D | Cat# MAB2457; RRID: AB_2159398 |
| Rat anti-PAX1 (1:100) | Abcam | Cat# ab252847 |
| Sheep anti-EBF2 (2.5 µg/mL) | R&D | Cat# AF7006; RRID: AB_10972102 |
| Goat anti-SOX17 (2.5 µg/mL) | R&D | Cat# AF1924; RRID: AB_355060 |
| Mouse anti-CD34 (1:200) | Bio-Techne | Cat# FAB7227P; RRID: AB_10973177 |
| Mouse anti-VE-cadherin (1:200) | CST | Cat# 2500; RRID: AB_10939118 |
| Mouse anti-N-cadherin (1:200) | BD | Cat# 610920; RRID: AB_2077527 |
| Goat anti-mClover (1:200) | St John’s Laboratory | Cat# STJ140283; RRID: AB_3717847 |
| Rabbit anti-RFP (mScarlet) (1:200) | Abcam | Cat# ab124754; RRID: AB_10971665 |
| Mouse anti-PAX6 (1:50) | DSHB | RRID: AB_528427 |
| Goat anti-NKX6-1 (2.5 µg/mL) | R&D | Cat# AF5857; RRID: AB_1857045 |
| Rabbit anti-OLIG2 (1:200) | CST | Cat# 65915T; RRID: AB_2936997 |
| Rabbit anti-Histone H3 (phospho Ser10) (1:200) | Abcam | Cat# ab47297; RRID: AB_880448 |
| Rabbit anti-CDX2 (1:200) | Thermo | Cat# A300-692A; RRID: AB_530295 |
| Mouse anti-FOXA2 (1:50) | DSHB | RRID: AB_528255 |
| Mouse anti-ZO-1 (1:200) | BD | Cat# 610966; RRID: AB_398279 |
| Mouse anti-FOXJ1 (1:200) | Thermo | Cat# 14-9965-82; RRID: AB_1548835 |
| Rabbit anti-ARL13B (1:200) | Proteintech | Cat# 17711-1-AP; RRID: AB_2060867 |
| Mouse anti-PAX7 (1:50) | DSHB | RRID: AB_528428 |
| Rabbit anti-BMP4 (1:200) | ABclonal | Cat# A11315; RRID: AB_2758503 |
| Rabbit anti-BMP7 (1:200) | ABclonal | Cat# A0697; RRID: AB_2757348 |
| Donkey anti-mouse IgG (H+L), Alexa 405 (1:200) | Abcam | Cat# ab175658; RRID: AB_2687445 |
| Donkey anti-mouse IgG (H+L), Alexa 488 (1:200) | Thermo | Cat# A21202; RRID: AB_141607 |
| Donkey anti-rat IgG (H+L), Alexa 488 (1:200) | Thermo | Cat# A21208; RRID: AB_2535794 |
| Donkey anti-rabbit IgG (H+L), Alexa 555 (1:200) | Thermo | Cat# A21206; RRID: AB_2535792 |
| Donkey anti-rabbit IgG (H+L), Alexa 647 (1:200) | Thermo | Cat# A31572; RRID: AB_162543 |
| Donkey anti-sheep IgG (H+L), Alexa 594 (1:200) | Thermo | Cat# A11016; RRID: AB_2534083 |
| Donkey anti-goat IgG (H+L), Alexa 555 (1:200) | Thermo | Cat# A31572; RRID: AB_162543 |
| Donkey anti-goat IgG (H+L), Alexa 647 (1:200) | Thermo | Cat# A21447; RRID: AB_2535864 |
| Chemicals, peptides, and recombinant proteins | | |
| DMEM/F-12 | Thermo or Servicebio | Cat# 21331020 or Cat# G4514 |
| Neurobasal | Thermo | Cat# 21103049 |
| GlutaMAX, 100x | Thermo | Cat# 35050061 |
| MEM NEAA, 100x | Thermo | Cat# 11140050 |
| Sodium pyruvate, 100x | Thermo | Cat# 11360039 |
| Antibiotic-Antimycotic, 100x | Thermo | Cat# 15240062 |
| Probumin | Sigma | Cat# 821001 |
| Transferrin (Holo) | Athens Research and Technology | Cat# 16-16-032001 |
| B27 supplement, 50x | Thermo | Cat# 17504001 |
| N2 supplement, 100x | Thermo | Cat# 17502048 |
| Trace Element A, 1000x | Corning | Cat# 25-021-CI |
| Trace Element B, 1000x | Corning | Cat# 25-022-CI |
| Trace Element C, 1000x | Corning | Cat# 25-023-CI |
| Ascorbic acid magnesium | TCI | Cat# A2521 |
| Accutase | Thermo | Cat# 00-4555-56 |
| TrypLE | Thermo | Cat# 12604-013 |
| Heregulin β1 | Qkine | Cat# QK045 |
| Activin A | Qkine | Cat# QK001 |
| IGF-1 LR3 | Qkine | Cat# QK041 |
| FGF2-G3 | Qkine | Cat# QK052 |
| FGF3 | MCE | Cat# HY-P700065 |
| FGF4 | MCE | Cat# HY-P700014 |
| FGF8b | Qkine | Cat# QK057 |
| FGF17 | MCE | Cat# HY-P700060 |
| WNT3A | R&D | Cat# 5036-WN |
| WNT5A | R&D | Cat# 645-WN |
| CER1 | MCE | Cat# HY-P7822 |
| SHH, high activity form | R&D | Cat# 8908-SH |
| BMP2 | Qkine | Cat# QK007 |
| BMP4 | R&D | Cat# 314-BP |
| BMP7 | R&D | Cat# 354-BP |
| All-trans Retinal (RAL) | Aladdin | Cat# R106320 |
| All-trans Retinoic acid (RA) | Aladdin | Cat# A122355 |
| Geltrex, growth factor reduced | Thermo | Cat# A1413302 |
| Matrigel, iPSC level | Mogengel | Cat# 827775 |
| Y-27632 | Aladdin | Cat# Y412681 |
| Chroman 1 | MCE | Cat# HY-15392 |
| Emricasan | MCE | Cat# HY-10396 |
| Polyamine supplement | Sigma | Cat# P8483 |
| Trans-ISRIB | Aladdin | Cat# I860930 |
| CHIR-99021 | MCE | Cat# HY10182 |
| SB-413542 | Aladdin | Cat# S125924 |
| LDN-193189 | Tocris | Cat# 1517128 |
| SAG-HCl | Aladdin | Cat# S872455 |
| SANT-1 | MCE | Cat# HY-100224 |
| Phalloidin Alexa Fluor 647 | CST | Cat# 8940S |
| Paraformaldehyde | Sigma | Cat# 158127 |
| DPBS | Servicebio | Cat# G4200 |
| SDS, 20% solution, RNase-free | Thermo | Cat# AM9820 |
| Triton X-100 | Sigma | Cat# T8787 |
| Tween-20 | Solarbio | Cat# T8220 |
| 20x SSC solution, RNase-free | Thermo | Cat# AM9770 |
| UltraPure water, DNase-/RNase-free | Thermo | Cat# 10977023 |
| DAPI | TCI | Cat# D5888 |
| OCT compound | Tissue-tek | Cat# 4583 |
| Isopentane | Macklin | Cat# I813377 |
| Critical commercial assays | | |
| E.Z.N.A. MicroElute Total RNA Kit | Omega Biotek | Cat# R6831-02 |
| iScript Reverse Transcription Supermix | Bio-Rad | Cat# 1708841 |
| Taq Pro HS Universal Probe Master Mix | Vazyme | Cat# QN113-01 |
| Duolink Blocking Solution, 1x | Sigma | Cat# DUO82007 |
| MAXbind^TM^ Staining Medium | Active motif | Cat# 15253 |
| MAXwash^TM^ Washing Medium | Active motif | Cat# 15254 |
| RNaseOUT^TM^ Recombinant Ribonuclease Inhibitor | Thermo | Cat# 10777019 |
| NeonTM Transfection System 100 µL Kit | Thermo | Cat# MPK10025 |
| ProLong Gold antifade mountant | Thermo | Cat# P36934 |
| HCR probes are listed in Table S1 | Molecular Instruments | Custom order |
| Deposited data | | |
| scRNA-seq data of hTLE.v1/2/3 | This manuscript | GSE314260 |
| Visium HD spatial scRNA-seq data of hTLE.v2,3 | This manuscript | GSE314260 |
| scRNA-seq data of mouse embryo development | Pijuan-Sala et al.^1^ | E-MTAB-6967 |
| scRNA-seq data of monkey embryo development | Zhai et al.^2^ | GSE193007 |
| scRNA-seq data of human CS7 embryo | Tyser et al.^3^ | E-MTAB-6967 |
| scRNA-seq data of human CS8 embryo | Xiao et al.^4^ | HRA005567 |
| scRNA-seq data of human CS10 embryo | Zeng et al.^5^ | GSE155121 |
| Experimental models: Cell lines | | |
| K3-iPSC (male) | Si-Tayeb et al.^6^ | N/A |
| WT H9-hESC (female) | WiCell | Cat# WAe009-A; RRID: CVCL_9773 |
| PAX3::mClover3/EBF2::mScarlet H9-hESC | This manuscript | N/A |
| NOTO::mClover3 H9-hESC | This manuscript | N/A |
| NKX1-2::mScarlet H9-hESC | This manuscript | N/A |
| MYF5::mClover3 H9-hESC | This manuscript | N/A |
| NOTO-LOF H9-hESC | This manuscript | N/A |
| VANGL1-LOF H9-hESC | This manuscript | N/A |
| VANGL2-LOF H9-hESC | This manuscript | N/A |
| VANGL1/2-LOF H9-hESC | This manuscript | N/A |
| Oligonucleotides | | |
| Oligos for gRNA plasmids are listed in Table S1 | BGI | Custom order |
| Taqman probes are listed in Table S1 | Thermo | N/A |
| Recombinant DNA | | |
| HDR donor arm vectors for CRISPR/Cas9 editing | BGI | Custom order |
| pRGS Surrogate Reporter | PNA Bio | Cat# RV01 |
| Software and algorithms | | |
| GraphPad Prism9 | GraphPad Inc. | https://www.graphpad.com/ |
| NIS-Elements AR (v5.41.01) | Nikon microscope | N/A |
| LAS X | Leica | N/A |
| BZ-X800 Analyzer | Keyence | N/A |
| FACSDiva software (v9.4) | BD | N/A |
| ImageJ | National Institutes of Health | https://imagej.nih.gov/ij/ |
| R | R Core Team | https://www.r-project.org/ |
| Python | Open source | https://www.python.org/ |
| cellranger (v9.0.1) | 10x Genomics | https://10xgenomics.com/support/software/cell-ranger/ |
| spaceranger (v3.1.3) | 10x Genomics | https://10xgenomics.com/support/software/space-ranger/ |
| Seurat (v4.3) | Satija Lab | https://satijalab.org/seurat/index.html |
| DoubletFinder (v2.0.4) | CS McGinnis et al.^7^ | https://github.com/chris-mcginnis-ucsf/DoubletFinder |
| celda (v1.18.2) | Wang et al.^8^ | https://github.com/campbio.celda |
| biomaRt (v2.62.0) | Bioconductor | https://bioconductor.org/packages/release/bioc/html/biomaRt.html |
| BBKNN | Polanski et al.^9^ | https://github.com/Teichlab/bbknn |
| harmony (v1.0.1) | Korsunsky et al.^10^ | https://github.com/immunogenomics/harmony |
| scvelo (v0.2.5) | Bergen et al.^11^ | https://github.com/theislab/scvelo |
| velocyto (v0.17.7) | La Manno et al.^12^ | https://velocyto.org |
| sceasy (v0.0.7) | Cellular Genomics Informatics | https://github.com/cellgeni/sceasy |
| Palantir (v1.3.3) | Setty et al.^13^ | https://github.com/dpeerlab/Palantir |
| SCENIC (v1.3.1) | Aibar et al.^14^ | https://github.com/aertslab/SCENIC |
| CellChat (v2) | Jin et al.^15^ | https://github.com/sqjin/CellChat |
| ClusterFoldSimilarity | González-Velasco et al.^16^ | https://github.com/OscarGVelasco/ClusterFoldSimilarity |
| Other | | |
| 96-well U-bottom plate, Sphera low-attachment surface | Thermo | Cat# 174929 |
| Ultra-low attachment 6-well plate | Thermo | Cat# 174929 |
| 16-well chamber slide | iBidi | Cat# 81811 |
| 100 µm cell strainer | SPL | Cat# SPL-39100 |
| QuantStudio 7 Pro Real-Time PCR System, 384-well | Applied Biosystems | Cat# A43183 |
| CO_2_ Resistant Shaker, Universal Platform | Thermo | Cat# 88881104 |
| Neon transfection system | Thermo | N/A |
| SU8010 cold-field Emission Scanning Electron Microscope | Hitachi | N/A |
| Q150T Automatic Coating System | Quorum Technologies | N/A |
| CM1950 Cryostat | Leica | N/A |
| SP8 inverted confocal microscope | Leica | N/A |
| BZ-X All-in-one inverted fluorescent microscope | Keyence | N/A |
| EVOS^TM^ M5000 imaging system | Thermo | N/A |
| Primovert inverted phase contrast microscope | ZEISS | N/A |
| Ti2-E inverted fluorescent microscope | Nikon | N/A |
| FACSAria^TM^ Fusion | BD | N/A |
| 10x Visium CytAssist | 10x Genomics | N/A |
| 10x Chromium iX Controller | 10x Genomics | N/A |
| C1000 Touch^TM^ Thermal cycler | Bio-Rad | N/A |
